# Quantitative assessment of NCLDV–host interactions predicted by co-occurrence analyses

**DOI:** 10.1101/2020.10.16.342030

**Authors:** Lingjie Meng, Hisashi Endo, Romain Blanc-Mathieu, Samuel Chaffron, Rodrigo Hernández-Velázquez, Hiroto Kaneko, Hiroyuki Ogata

## Abstract

Nucleocytoplasmic DNA viruses (NCLDVs) are highly diverse and abundant in marine environments. However, knowledge of their hosts is limited because only a few NCLDVs have been isolated so far. Taking advantage of the recent large-scale marine metagenomics census, *in silico* host prediction approaches are expected to fill the gap and further expand our knowledge of virus–host relationships for unknown NCLDVs. In this study, we built co-occurrence networks of NCLDVs and eukaryotic taxa to predict virus–host interactions using *Tara* Oceans sequencing data. Using the positive likelihood ratio to assess the performance of host prediction for NCLDVs, we benchmarked several co-occurrence approaches and demonstrated an increase in the odds ratio of predicting true positive relationships four-fold compared with random host predictions. To further refine host predictions from high-dimensional co-occurrence networks, we developed a phylogeny-informed filtering method, Taxon Interaction Mapper, and showed it further improved the prediction performance by twelve-fold. Finally, we inferred virophage – NCLDV networks to corroborate that co-occurrence approaches are effective for predicting interacting partners of NCLDVs in marine environments.

**Importance:** NCLDVs can infect a wide range of eukaryotes although their life cycle is less dependent on hosts compared with other viruses. However, our understanding of NCLDV– host systems is highly limited because few of these viruses have been isolated so far. Co-occurrence information has been assumed to be useful to predict virus–host interactions. In this study, we quantitatively show the effectiveness of co-occurrence inference for NCLDV host prediction. We also improve the prediction performance with a phylogeny-guided method, which leads to a concise list of candidate host lineages for three NCLDV families. Our results underpin the usage of co-occurrence approach for metagenomic exploration of the ecology of this diverse group of viruses.

## Introduction

Nucleocytoplasmic large DNA viruses (NCLDVs) represent a group of double-stranded DNA viruses that belong to the viral phylum *Nucleocytoviricota* (<u>Virus Taxonomy: 2019 Release</u>), which was previously referred to as Megavirales (1, 2). NCLDVs usually possess diverse gene repertoires (74 to more than 2,000 proteins), large genomes (45 kb to 2.5 Mb), and outsized virions (80 nm to 1.5 µm) (3–5). NCLDVs have high functional autonomy and encode components of replication, transcription, and translation systems (3). Recently, a virus that belongs to a new family of NCLDVs called “Medusaviridae” was found to encode five types of histones (6). The existence of metabolically active viral factories and infectious virophages also indicates that the life cycle of NCLDVs is less dependent on host cells than other viruses (7, 8). To further understand what makes these giant viruses more independent than other viruses, a first crucial step is to identify their hosts — “Who infects whom?”.

NCLDVs are known to infect a broad range of eukaryotes, from unicellular eukaryotes and macroalgae to animals (9). Amoebae are frequently used hosts in co-culture to isolate large NCLDVs (10). However, there is growing evidence, especially in marine systems, that NCLDVs can infect many phytoplankton groups, such as Pelagophyceae, Mamiellophyceae, Dinophyceae, and Haptophyte (11–13). Several other non-photosynthetic eukaryotic lineages, such as Bicoecea and Choanoflagellatea, were also reported as experimentally identified NCLDV hosts in marine environments (14, 15). Small to large marine organisms, including invertebrates and vertebrates, are infected by viruses that belong to the NCLDV family *Iridoviridae* (16, 17). Together these studies indicate ubiquitous infectious relationships between NCLDVs and a wide range of marine eukaryotes. However, our understanding of NCLDV–host systems is very limited because few viruses have been isolated so far.

The number of viruses and hosts isolated in the laboratory represents a very small fraction of existing interactions in the ocean. Indeed, NCLDVs have been found to be highly diverse and abundant based on omics data (18, 19). In only a few liters of coastal seawater, more than 5,000 *Mimiviridae* species were detected; by comparison, only 20 *Mimiviridae* with known hosts have been well investigated (20). Global marine metagenomic data have revealed that the richness and phylogenetic diversity of NCLDVs are even higher than those of an entire prokaryotic domain (21). From biogeographical evidence, it is clear that these viruses are prevalent in the marine environment but have a heterogeneous community structure across sizes, depths, and biomes (22). Marine metatranscriptomic data have also shown that NCLDVs are active everywhere in sunlit oceans and may infect hosts from small piconanoplankton (0.8–5 µm) to large mesoplankton (180–2000 µm) (23).

Previous studies also demonstrated that NCLDVs have the potential to infect a greater diversity of hosts than known to date through gene transfer analyses (24, 25). NCLDVs might have started coevolving with eukaryotes even before the last eukaryotic common ancestor (LECA) (26). A recent study supported this hypothesis by showing that some NCLDVs encode viractins (actin-related genes in viruses), which could have been acquired from proto-eukaryotes and possibly reintroduced in the pre-LECA eukaryotic lineage (27). Together, these findings underline a lack of knowledge about NCLDV biology and host diversity. Therefore, more effort is needed to identify hosts to elucidate the poorly known virus–host relationships and the largely unknown NCLDV world.

Substantial effort has been made to reveal interactions between NCLDVs and their putative hosts. Apart from the co-culture method, other alternative methods, including high-throughput cell sorting, are also being used (10, 15). Metagenomics, which is particularly useful to assess a large fraction of ecosystem diversity, has been increasingly used to investigate NCLDVs host range. Comparative genomics analyses, such as identification of horizontal gene transfer (HGT) predictions, have also performed well for NCLDV host prediction (24, 25).

Abundance-based co-occurrence analyses have been used for host prediction and are supposed to be effective because viruses can only thrive in an environment where their hosts exist (18, 28). In addition to virus–host relationships, co-occurrence networks have been used to predict the association between NCLDVs and their “parasites” (virophages) (29). However, the co-occurrence-based prediction is also controversial for viral host prediction since the abundance dynamics of viruses and their hosts (e.g., *Emiliania huxleyi* and *Heterosigma akashiwo* viruses) are sometimes not concordant (30, 31). Usually, validation with known virus–host relationships or corroboration with genomic evidence (e.g., HGT) is used to assess network-based predictions (18, 28). However, the effectiveness of previous and novel co-occurrence network methods has never been quantitatively tested for NCLDV host prediction. The current lack of quantitative assessment hinders the widespread use of this approach. Therefore, dedicated methods are needed to test the accuracy of NCLDV host prediction with co-occurrence networks and to improve the performance of co-occurrence-based predictions.

The *Tara* Oceans expedition is a global-scale survey on marine ecosystems that expands our knowledge of microbial diversity, organismal interactions, and ecological drivers of community structure (32). The present study used *Tara* Oceans metagenomic and metabarcoding datasets to predict virus–host relationships between NCLDVs and eukaryotes by constructing co-occurrence networks using different methods. To quantitatively assess the performance of network-based host prediction, we employed the positive likelihood ratio (LR+) using reference data for known NCLDV–host relationships. We developed a phylogeny-based enrichment analysis approach, Taxon Interaction Mapper (TIM), to enhance the performance in detecting positive signals in the intricate inferred networks. TIM has previously been used in host predictions for DNA and RNA viruses (33), but without a quantitative assessment on its effectiveness. In this study, we assessed the performance of TIM as a filter of co-occurrence networks. We examined NCLDV–virophage networks, which further justify the use of co-occurrence and filtering approaches to identify NCLDV interaction partners.

## Results

### NCLDV–eukaryote co-occurrence networks

From five datasets that corresponded to five size fractions (Fig. S1), we generated five co-occurrence networks on a global scale (Fig. 1, S2A). Altogether, these networks were composed of 20,148 V9 and 5,234 *polB* OTUs (nodes) and 47,978 *polB*–V9 associations (edges). Out of these associations, 47,296 had positive weights, and 682 had negative weights (Fig. 2A). The associations that involved the family *Mimiviridae* were numerically dominant (*n* = 36,830) among the different NCLDV families. The second largest family was *Phycodnaviridae*, with 5,521 edges involving eukaryotes. No other family had more than 2,000 associations with eukaryotes. *Marseilleviridae*, forming the least associations in the networks, had 132 edges with eukaryotes. Taxonomic annotation of eukaryotic OTUs indicated that Alveolata, Opisthokonta, Rhizaria, and Stramenopiles were the major four eukaryotic groups connected to NCLDVs (with 21,167, 9,179, 6,521, and 5,327 edges, respectively). Three of these eukaryotic groups belong to the SAR supergroup (i.e., Stramenopiles, Alveolata, and Rhizaria), which represented 68.81% of the total associations. Regarding the pairs between viral families and eukaryotic lineages, *Mimiviridae* and Alveolata showed the largest number of edges (*n* = 16,548). Besides NCLDV–eukaryote associations, we detected 57,495 *polB–polB* associations and 234,448 V9–V9 associations (Fig. S2B). We also included environmental parameters in the network inference and identified 25 pairs of associations between environmental parameters and *polB* OTUs (Table S1).

**Figure 1.**
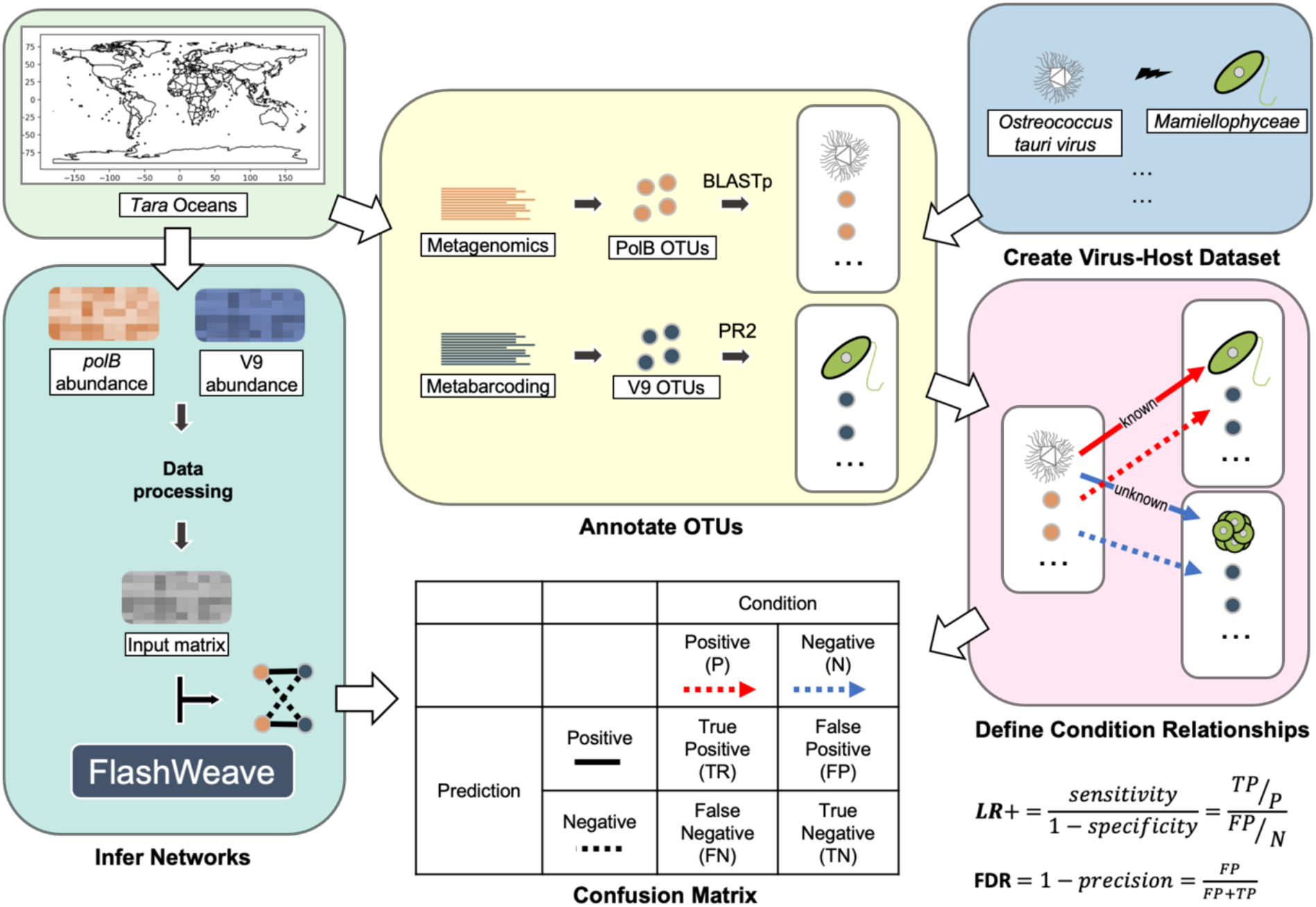
Overall workflow for inferring co-occurrence networks and quantitative assessment. This figure shows how the input data (*Tara* Oceans metagenomics and metabarcoding data) were used in this study. The definition of the confusion matrix for quantitative assessment is shown in the table. The LR+ and FDR equations are given at the lower right corner of the plot.

**Figure 2.**
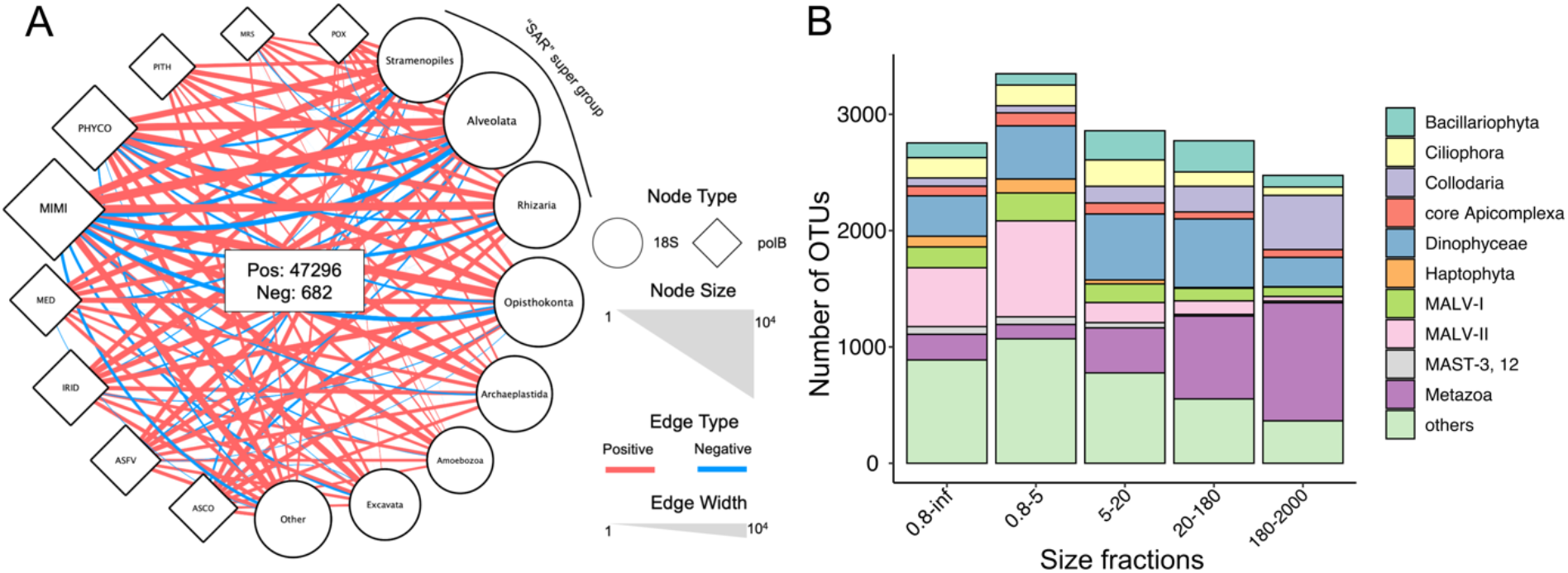
*polB*–V9 co-occurrence network. We performed co-occurrence analysis at the OTU level and constructed the network with pooled *polB* –V9 associations from five size fraction networks (A). To better display co-occurrence patterns, PolB OTUs were grouped at the family or family-like level, and V9 OTUs were grouped using annotation at high taxonomic ranks. The size of each node indicates the number of OTUs that belong to the group, and the width of each edge indicates the number of associations between two connected groups. Associations with positive weight are shown in red and negative associations are shown in blue. (B) Number of associations connected to NCLDVs for each major eukaryotic lineage in five size fractions. The top 10 lineages were retained, and other lineages were omitted and shown as “others.” Size fractions are presented in μm.

The number of NCLDV–eukaryote associations generally decreased with enlarging size fraction (Fig. S2A). The largest number of *polB–*V9 associations were found in the 0.8– 5-μm fraction (*n* = 10,647). Correspondingly, the eukaryotic community in 0.8–5-μm fraction had the greatest diversity (Fig. S3). However, the 0.8–inf-μm size fraction network was the largest (*n* = 10,477) for edges with positive weights. With the annotation of major lineages, the eukaryotic community compositions in the networks varied across different size fractions (Fig. 2B). In the smallest size fraction (0.8–5 μm) and the large range size fraction (0.8–inf μm), Marine Alveolate Group II was the eukaryotic lineage with the largest number of associations with NCLDVs (21.39% and 19.98%, respectively). Dinophyceae was the second largest group connected to NCLDVs in these two size fractions and showed the largest number of connections with NCLDVs in the 5–20-μm size fraction network (22.22% of total interactions). The viral associations with Metazoa and Collodaria increased with increasing size fractions. In the largest 180–2000-μm size fraction network, Metazoa contributed 39.31% of the total *polB*–V9 edges.

We calculated the degree of nodes (number of connected edges) for each NCLDV *polB* OTU (Fig. 3A, B). Naturally, the average degree of positive associations per *polB* was higher than negative edges in all size fractions and decreased along with increasing size fractions (2.69, 2.40, 2.25, and 2.10 from 0.8–5 μm to 180–2000 μm, and 2.76 for 0.8–inf). Most of the *polB* nodes had more than one positive association (Fig. 3A). Together with the taxonomic annotation of nodes, *polB*–V9 associations in the networks generated with the *Tara* Oceans data revealed their high dimensionality and complexity.

**Figure 3.**
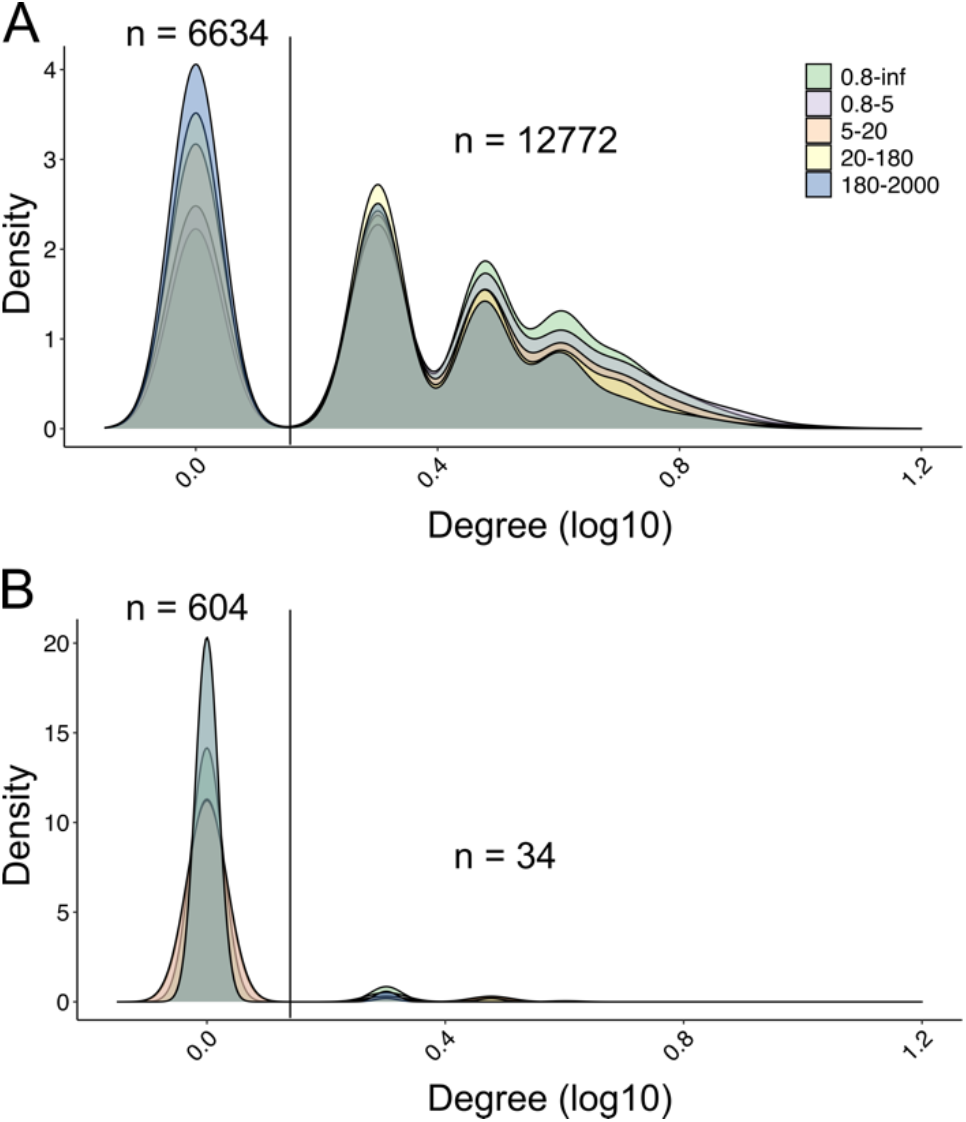
Density plots for the degree of NCLDV nodes in co-occurrence networks. The degree of an NCLDV node is given by the associations between this node and eukaryotes in the networks. The amount of NCLDV nodes are given on the top of the density values. (A) Positive degree (number of positive associations per node) for NCLDV nodes in five size fraction networks. (B) Negative degree (number of negative associations per node) for NCLDV nodes in five size fraction networks. Size fractions are presented in μm. NCLDV nodes with degree = 1 and degree > 1 are separated using a vertical line, and the number of nodes is given.

### Network validation

We quantitatively assessed the performance of predicting *polB*–V9 associations using the positive likelihood ratio (LR+) (Fig. 1). By defining groups of metagenomic PolBs as described in the Materials and Methods, 932 OTUs were recruited in the validation, and these sequences contributed 6191 *polB*–V9 associations in the FlashWeave networks (Fig. S4). To obtain an overall performance, we assessed the pooled associations (by removing redundancy) from the five co-occurrence networks. LR+ was separately calculated for edges with positive and negative weights because they can represent different infectious patterns. As shown in Fig. 4A, the LR+ of host prediction for positive associations was higher than 1 (LR+ = 1 indicates no change in the likelihood of the condition). The LR+ generally increased with the cut-off for FlashWeave weights, which indicated that condition positive cases are enriched in the edges with higher weights. This result demonstrated that the co-occurrence-based host prediction of NCLDVs outperformed random prediction (i.e., random inference of virus–host pairs). In high-weight regions: 1) weight > 0.6, the LR+ of associations was higher than 10; 2) weight > 0.4, the LR+ was roughly higher than 4. Nonetheless, the false discovery rate (FDR) was high (Fig. S5A), which indicated that the predictions contained numerous virus–host edges that were not considered condition positive. FDR was 91.67% and 96.34% when weight is greater than 0.6 and 0.4, respectively. An assessment of the host prediction for negative weight associations was also carried out. There were no known NCLDV–host pairs found in the negative networks (Fig. S5B). The analysis of the remaining part of our study was thus conducted for positive associations.

**Figure 4.**
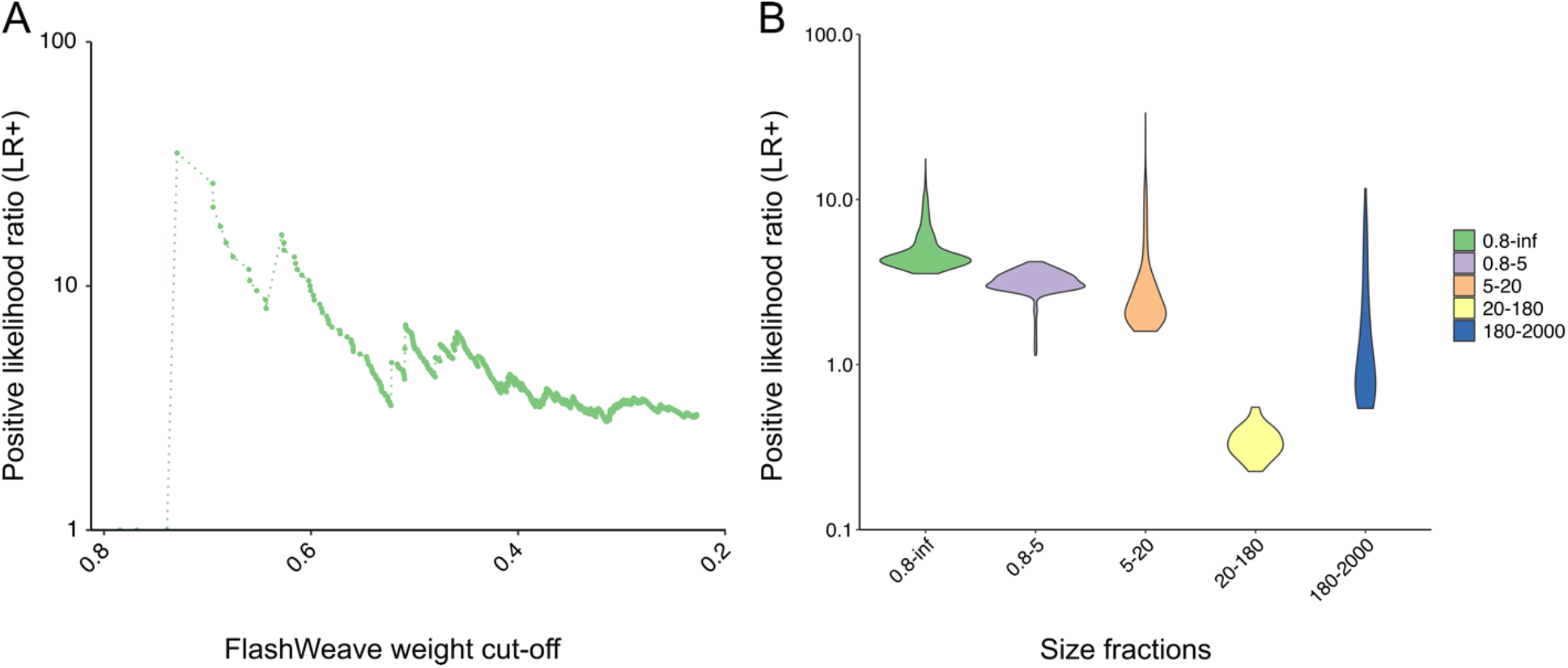
Positive likelihood ratios (LR+) in the NCLDV virus–host validation. (A) General performance of co-occurrence networks is shown with the LR+ calculated with associations pooled from five size fractions networks. To show the relationship between LR+ and FlashWeave association weight, the LR+ values are plotted along with the association weight. (B) Performance of each size fraction network is shown with the violin plot by ggplot2 with a bandwidth of 2. Size fractions are presented in μm.

Comparing the performance between different size fractions indicated that the networks of small size fractions (including the 0.8–inf-μm size fraction) performed better in predicting the NCLDV–host relationships (Fig. 4B, S6). The 0.8–inf-μm size fraction had the highest average LR+ out of the five size fractions (LR+ = 4.97). The LR+ of small size fractions was generally higher than that of large size fractions, but there were exceptions between 180–2000 and 20–180 μm. The LR+ of the associations in the 0.8–inf-μm, 0.8–5-μm and 5–20-μm was greater than 1. Different from the average results, when the weight is greater than 0.8, the associations of 5–20-μm size fraction had the best performance in terms of both LR+ and FDR (Fig. S6 A, B).

We also compared abundance filtration strategies using Flashweave-S (sensitive model) and FlashWeave-HE (heterogeneous model) but did not find a consistent pattern in prediction performance (Fig. S7). The networks from the Q1 filtration strategy performed best using Flashweave-S, but Q1 (lower quartile) filtration was not better than Q2 (middle quartile) for Flashweave-HE inferred networks. Flashweave-S had a better performance than HE model with any filtration strategy. Finally, we compared the performance of networks inferred by all three methods: FlashWeave-S, FastSpar and Spearman. Three methods generated a comparable number of positive associations, but FlashWeave-S made the largest number of true positive predictions (Fig. S8A). Noteworthy, the LR+ of these three methods were all larger than 1, however, FlashWeave-S and Spearman performed better than FastSpar (Fig. S8B).

### Assessment of host prediction improvement

Then we used the newly developed phylogeny-guided host prediction tool, TIM, to filter *polB*–V9 associations, which is based on the assumption that evolutionarily related viruses tend to infect evolutionarily related hosts (see Materials and Methods). We identified 24 eukaryotic taxonomic groups specifically associated with NCLDVs (Fig. S9). To compare the performance of the TIM results with the above raw FlashWeave results, we converted the three primary eukaryotic taxonomic ranks to their associated major lineages (Table S2), and the associations were plotted as a network (Fig. 5A). This network showed that three out of nine NCLDV families (*Mimiviridae, Phycodnaviridae*, and *Iridoviridae*) had enriched connections in specific eukaryotic lineages. Among the network edges, known virus–host pairs were found, such as Haptophyta–*Mimiviridae*, Mamiellophyceae–*Phycodnaviridae*, and Metazoa–*Iridoviridae*. The associations in the TIM-filtered results showed a sharp improvement in performance from the original result with and without an edge weight cut-off. The average LR+ of TIM-enriched associations was 42.22, which was higher than the raw FlashWeave associations without a weight cut-off (3.43), with a weight cut-off of 0.4 (5.20), and with a cut-off at 0.668 (14.23) (Fig. 5B, S9A). The FDR dropped from 0.97 (no cut-off) and 0.95 (weight cut-off of 0.4) to 0.74 (Fig. 5C).

**Figure 5.**
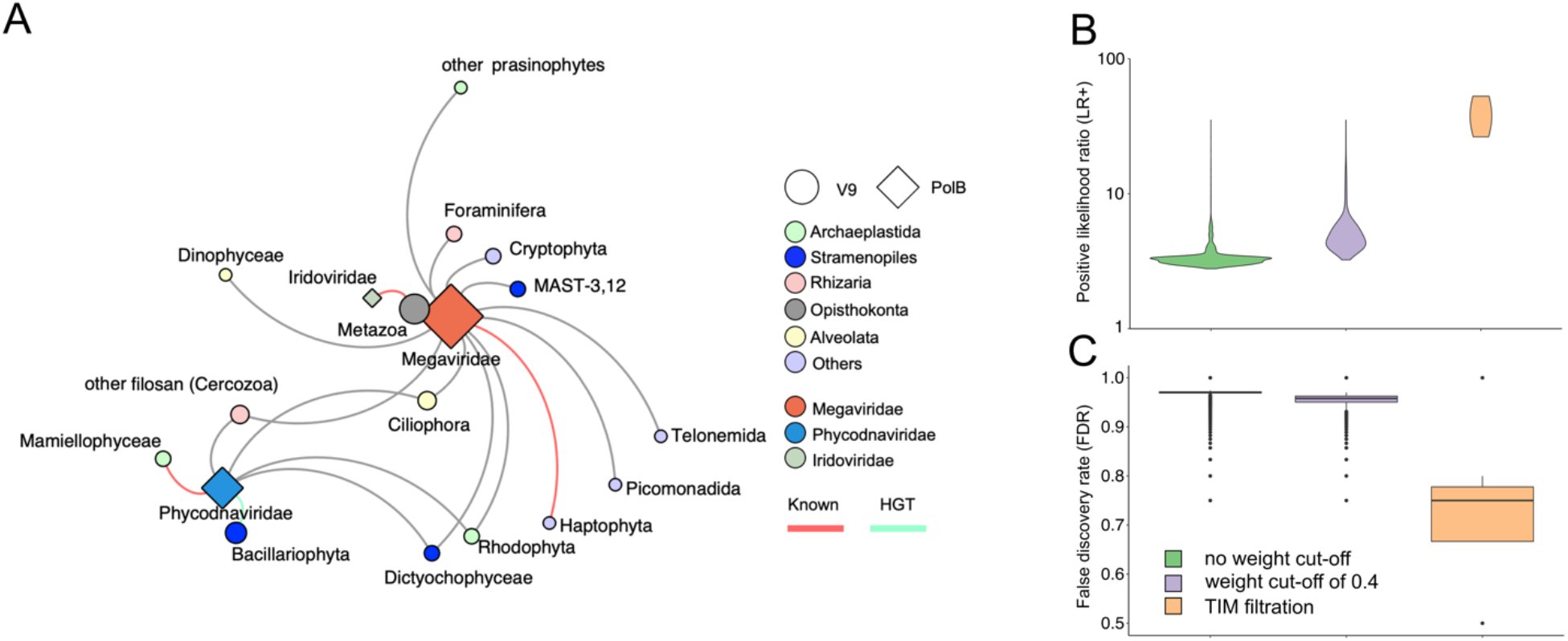
Prediction of NCLDV virus–host relationships with TIM. (A) Undirected network that shows the relationships between NCLDVs and eukaryotes after TIM filtration. The size of each node indicates the number of predicted interactions of this group. The weight of network edges as defined by the number of tree nodes enriched in each viral family subtree to specific eukaryotic major lineages in the TIM analysis. Known virus–host relationships are highlighted in red, and the pairs found to have horizontal gene transfer are highlighted in yellow (1). (B) Performance of networks on NCLDV host prediction for original FlashWeave results without a weight cut-off, weight cut-off > 0.4, and TIM filtration, plotted by ggplot2 with a bandwidth of 2. (C) FDR of networks for NCLDV host prediction with the original FlashWeave results without a weight cut-off, weight cut-off > 0.4, and TIM filtration.

From the network, diverse putative hosts (13 lineages) emerged for *Mimiviridae*, including algae, protozoans, and metazoans. Metazoa had the most enriched nodes connected to *Mimiviridae*; additionally, MAST-3,12, Cryptophyta, Foraminifera, and Ciliophora had strong relationships with *Mimiviridae*. For *Phycodnaviridae*, there were six eukaryotic lineages retained after TIM filtration. Among these, Bacillariophyta, “other filosan (part of filosan Cercozoa)”, and Mamiellophyceae had comparatively strong associations. Moreover, Rhodophyta, Ciliophora, and Dictyochophyceae had links to both *Mimiviridae* and *Phycodnaviridae*. There was also a connection between *Iridoviridae* and Metazoa.

### Associations between virophages and NCLDVs

Using 6,818 NCLDV *polB* OTUs and 195 virophage major capsid proteins (MCPs), we identified 535 FlashWeave associations (196 and 339 for pico- and femto-size fractions, respectively) (Fig. 6A). Most of the associations had positive weights (*n* = 490), whereas some had negative weights (*n* = 45). The average number of associations per virophage MCP was different in two size fractions: 3.2 in femto- and 5.6 in pico-size fractions. The network revealed that *Mimiviridae* had the largest number of virophage associations in both size fractions. We also detected 84 positive associations between virophages and *Phycodnaviridae*.

**Figure 6.**
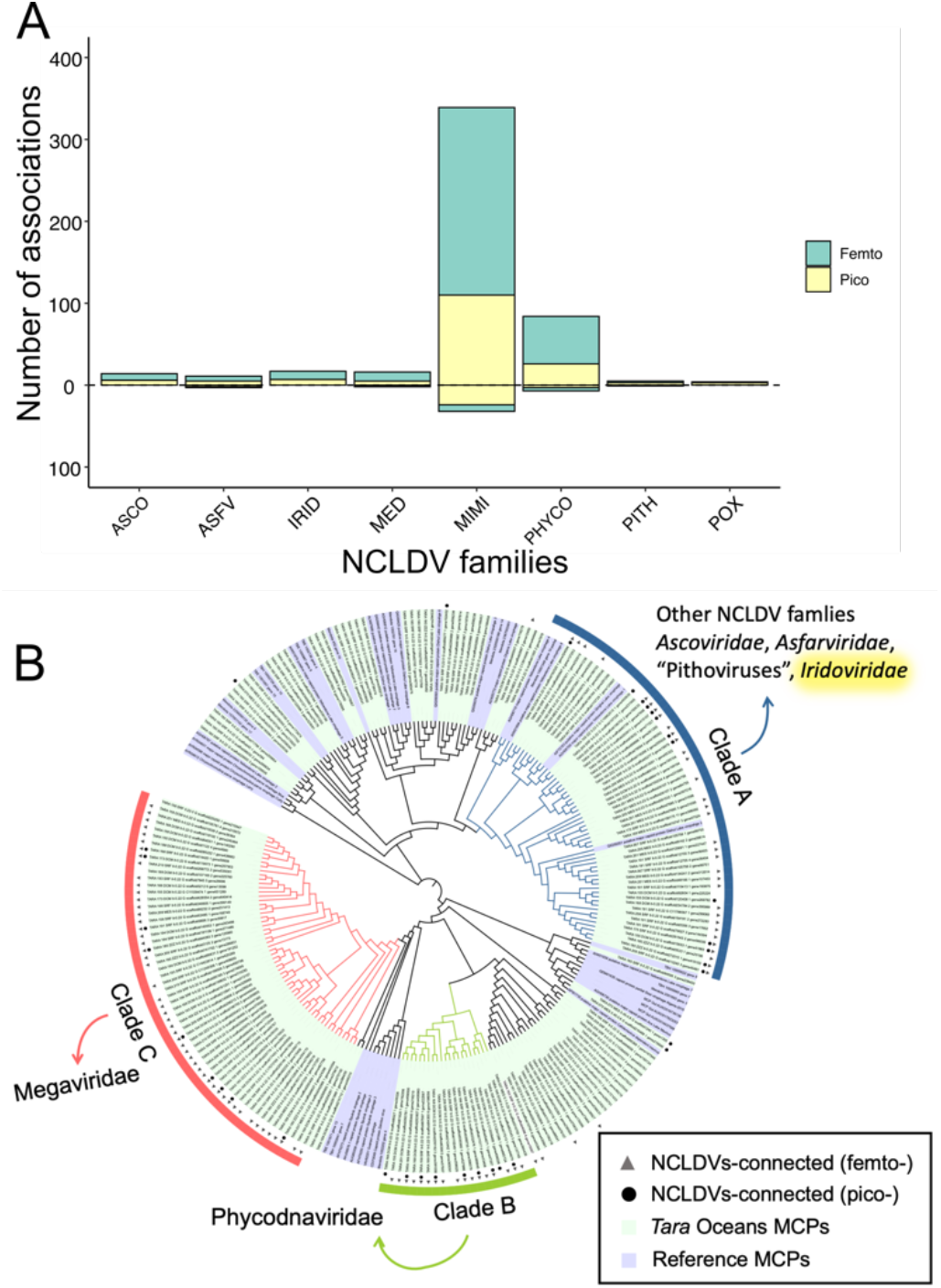
Associations between virophages and NCLDVs. (A) Number of associations with virophages is shown for seven NCLDV families and two unclassified groups, “Medusaviruses” and “Pithoviruses.” Associations in the femto-size fraction network are shown in yellow, and in the pico-size fraction network are shown in green. The number of positive associations is above the zero axis, and the number of negative associations is below the zero axis. (B) Phylogenetic tree was constructed from 195 environmental virophages and 47 reference MCP sequences. The outside layer indicates three major virophage clades. The inner two layers indicate that the virophage OTUs have at least one association with NCLDVs in femto- or pico-size fractions networks.

The phylogenetic tree defined three main virophage clades, and they were all found to have many connections to NCLDVs. To investigate significant relationships, Fisher’s exact test was performed between virophage clades and NCLDV families. Families other than *Phycodnaviridae* and *Mimiviridae* did not show significant associations. Therefore, we made a group, “Other NCLDVs,” to include all families except *Phycodnaviridae* and *Mimiviridae*. First, we only used FlashWeave results with a weight > 0.4, as previous results showed that a FlashWeave weight of 0.4 is a suitable cut-off that produced moderate performance (Fig. 3A). From the femto-size fraction network, we found two significantly enriched connections (Fig. 6B): one was between virophage group C and *Mimiviridae* (*p* = 0.0022) and the other was between group A and “Other NCLDVs” (*p* = 0.0439). Another significantly enriched relationship between virophage group B and *Phycodnaviridae* (*p* = 0.0410) was found in pico-size fractions when we used all associations without edge weight cut-off.

Finally, we examined HGTs of virophage MCPs in NCLDV genomes. We found two HGTs of virophage MCPs; both showed links between clade A and *Iridoviridae* (Table S3). This result was consistent with the Fisher’s exact test result, which revealed a connection between virophage clade A and “Other NCLDVs” including *Iridoviridae*.

## Discussion

NCLDVs can infect a wide range of eukaryotes, from unicellular to multicellular organisms (34). However, we are still far from a comprehensive knowledge of their hosts because few have been isolated so far. Therefore, better host prediction algorithms are needed to understand the ecological functions and evolutionary significance of NCLDVs. To make these predictions, we constructed global ocean co-occurrence networks based on the marine metagenome and metabarcoding datasets from 85 stations of the *Tara* Oceans expedition, which cover all major oceanic provinces across an extensive latitudinal gradient from pole to pole. The edges (associations) between *polB* and V9 nodes (OTUs) in the networks were generated using FlashWeave. The networks were particularly dense (Fig. 2A, S2A), thus suggesting that NCLDVs interact with numerous eukaryotes in the ocean. This was expected given the high abundance and diversity of NCLDVs in marine environments (18, 21) and the identification of HGT between these viruses and diverse eukaryotic lineages (24). The networks were dominated by the *Mimiviridae* nodes, which is consistent with previous reports that *Mimiviridae* is the most abundant and has the widest array of transcribed genes out of NCLDV families in marine environments (22, 23). *Mimiviridae* was known to infect amoebae, algae, and stramenopiles (3). In our study, these three eukaryotic groups were all found to have numerous associations with *Mimiviridae. Phycodnaviridae* has been known to infect many species of aquatic organisms, such as *Emiliania huxleyi* (Haptophyta), *Ectocarpus siliculosus* (Phaeophyceae), *Chlorella heliozoae* (Trebouxiophyceae), and *Ostreococcus tauri* (Mamiellophyceae) (35–37). Correspondingly, plenty of associations of *Phycodnaviridae* were found in the co-occurrence networks. For the eukaryotic nodes, all high taxonomic rank groups, including the SAR supergroup (i.e., Stramenopiles, Alveolata, Rhizaria), Opisthokonta, Archaeplastida, Amoebozoa, Excavata, and other eukaryotes, have associations with NCLDVs. Among these groups, the SAR supergroup contributed the most (∼68%) *polB*–V9 associations. However, this is still lower than in other microbial co-occurrence analyses; for example, a previous study showed SAR supergroup dominated ∼92% of the total aquatic microbial associations (38). A substantial proportion (∼32%) of NCLDV–eukaryote interactions were from non-SAR groups, which covered the known NCLDV host range, such as Archaeplastida and Haptophyta.

However, it is difficult to accurately predict NCLDV hosts from constructed networks because of the high degree of associations per *polB* OTU (Fig. 3A). One node connected to multiple edges was expected in the co-occurrence analysis. In previous NCLDV host prediction studies, additional processing was performed to filter the high dimensional associations to predict the meaningful interactions, such as weight cut-off or a combination of different co-occurrence network inference methods (18, 29). Moreover, no previous study quantitively assessed the performance of co-occurrence networks when predicting NCLDV– host relationships. Qualitatively identifying known pairs and detecting HGT (without validation) have been commonly used to assess prediction reliability (18, 28). Therefore, we aimed to 1) quantitatively assess the performance of co-occurrence-based host prediction for NCLDVs and 2) improve the prediction results using filtering methods.

In a previous study of bacteriophage host prediction, ROC curves were used as an assessment metric to compare different prediction methods (39). However, the number of known virus–host pairs of NCLDVs is not sufficient to generate a dataset for ROC assessment. Therefore, in this study, we carried out an alternative method, the LR+, to assess the performance. LR+ is calculated with two relative values, sensitivity and specificity (Fig. 1). The LR+ of co-occurrence-based host predictions for positive associations was higher than 1 and increased along with increasing cut-off values for the edge weights (Fig. 4A), which demonstrated that positive prediction results were more likely to be true positives than those based on random prediction. In high-weight regions (> 0.6 and > 0.4), LR+ values were larger than 10 and 4, respectively. These LR+ values indicate that FlashWeave can increase the probability of predicting true positives (40). However, both the true positive rate (sensitivity, < 18.9%) and false positive rate (< 6.37%) were very low (Fig. S5C). These low rates were from FlashWeave with a cut-off of alpha < 0.01, which excluded a large proportion of the *polB*–V9 pairs from the results. So only about 4000 predictions could be validated from a set of 6191 *polB*–V9 FlashWeave associations in this study (Fig. S5D, as described in the Materials and Methods). The FDR of co-occurrence was even higher than 90% (Fig. S5A). Such a high FDR in co-occurrence networks demonstrates that condition positive connections (i.e., known interactions) are embedded in an immense pool of condition negative connections. However, these negative signals can correspond to either unidentified (i.e., currently unknown true interactions), indirect, or false relationships.

We also found that true positive predictions only existed in positive weight associations, whereas negative weight associations did not contribute to NCLDV–host detection (Fig. S5B). This result indicates that the abundance dynamics of NCLDVs and their potential hosts were positively correlated with each other in the analyzed samples, which were collected at a global scale; this might be because NCLDVs detected in the dataset were active viruses that replicate locally in their hosts. Similar results were obtained in other co-occurrence-based host prediction studies (28, 41). However, several experimental studies showed that the abundance dynamics of NCLDVs and hosts showed a delay in time (30, 31). It is possible that the global-scale samples did not have sufficiently high resolution to detect negative correlations (or correlations with a time delay) due to lack of time-resolution (42). Therefore, further studies, especially those that focus on a high temporal resolution, are needed to better understand the detailed dynamics of virus–host associations and the capacity of co-occurrence-based methods for host prediction.

The networks of different size fractions showed different performance patterns in predicting NCLDV–host relationships (Fig. 4B). This pattern is not dependent on the diversity of eukaryotic communities (Fig. S3A, B). Generally, small-sized fractions (0.8–5-µm and 5–20-µm) networks performed better than large-sized fractions (20-180-µm and 180-2000-µm) networks. This result is not dependent on the diversity of eukaryotic communities. To date, most of the known NCLDV hosts are small, such as the genera *Micromonas, Aureococcus*, and *Ostreococcus* are within the range of 0.8–5-µm, and *Prymnesium, Heterosigma*, and *Heterocapsa* are within the range of 5–20-µm. Because of this, our assessment method might be biased toward small size fractions as smaller organisms tend to be more abundant in the environment (43). However, it is also possible that NCLDV infections are more prevalent in smaller size fractions. Notably, the 0.8–inf-µm size fraction network, which covered all four individual size fractions, performed best. This might be because NCLDVs can infect not only small hosts but also hosts from a broad size range.

Trimming of low-abundance OTUs was recommended to improve the prediction of true interactions and was often used in co-occurrence studies (44, 45). In our study, however, we did not achieve such performance improvement by treating input abundance data with a rigorous filtration (upper quartile) (Fig. S7). This result might be because the true positive and false positive rates defined in this study were too low; therefore, the validation may not be sufficiently sensitive to reflect the change between different abundance trimming strategies. However, it is also possible that low-abundance NCLDV OTUs are indeed network participants, as was demonstrated in a study showing that rare cyanobacterial species might play fundamental roles in blooming (46). Our result also revealed that FlashWeave-S was better than FlashWeave-HE at predicting NCLDV–host interactions (Fig. S7). The difference between FlashWeave-HE and FlashWeave-S is that HE mode can remove structural zeros during network inference. Structural zero is a typical property of heterogeneous datasets, like *Tara* Oceans datasets, and may lead to false-positive edges (47). Conversely, our results suggested that retaining structural zeros did not negatively influence the result, which indicates that the “presence–absence” pattern is as informative as the “more–less” pattern when identifying NCLDV–host relationships. This result is consistent with a previous “K-r-strategist” hypothesis: some NCLDVs, like mimiviruses, are K-strategists that decay slowly and can form stable associations with their hosts (48, 49). A recent report supported these non-”boom and bust” dynamics of prasinoviruses and their hosts with an experiment-based mathematical model (50). Overall, our results support co-occurrence networks as a useful method for predicting NCLDV–host interactions in marine metagenomes, and likelihood ratios as useful quantitative metrics for assessing the performance of co-occurrence analysis for viral host predictions.

Although the results generated by FlashWeave already improved the accuracy of predictions, the condition positive interactions were still embedded in many noise edges, as shown by a very high FDR (Fig. S5A, S6B). To overcome this situation, we developed TIM to reduce the high dimension of associations and improve NCLDV host prediction (33). The results showed that NCLDVs had enriched connections with 15 major eukaryotic lineages, which included 24 taxonomic groups in three different ranks (order, class, and phylum) (Table S2) (Fig. 5A, S7). Using the LR+ as a prediction diagnostic metric, NCLDV host prediction improved 12-fold with TIM filtration (Fig. 5B). FDR dropped below 23% after TIM treatment (Fig. 5C). In TIM-enriched connections, some are known NCLDV–host pairs, such as *Phycodnaviridae* and Mamiellophyceae, *Mimiviridae* and Haptophyta, and *Iridoviridae* and Metazoa. Some other studies revealed that *Mimiviridae* could exclusively infect diverse putative hosts (24, 51). Our results support the assumption that *Mimiviridae* has connections with 13 eukaryotic lineages out of 15 total lineages. Among these lineages, *Mimiviridae* had the most numerous links to Metazoa. Some mimiviruses (namaoviruses) are known to infect freshwater sturgeon, *Acipenser fulvescens* (52). Metazoans are presumed to be susceptible to mimiviruses, because the choanoflagellates, a group of eukaryotes that is phylogenetically close to metazoans, were recently identified to be the host of a species of *Mimiviridae* (15). Moreover, the TIM result revealed that *Phycodnaviridae* is closely connected to Bacillariophyta, which consists of three NCBI taxonomic groups: Thalassiophysales, Cymbellales, and Bacillariophyceae. Thalassiophysales was shown to have many HGT candidates with a large range of NCLDVs, and Bacillariophyceae also has a significant HGT candidate with phaeoviruses (24). Although Dictyochophyceae itself has not been proven to be a phycodnavirus host, its sister group Pelagophyceae was experimentally identified as an AaV host (53). Additionally, it is interesting to note the connection between Metazoa (Calanoida) and *Iridoviridae*. Calanoida is an order of arthropods commonly found as zooplankton; most of the sizes are 500–2000 µm. The viruses of the family *Iridoviridae* infect many Arthropod species, including insects and crustaceans (17).

Furthermore, we also inferred associations between virophages and NCLDVs. To date, all isolated virophages are only known to infect *Mimiviridae* (54). As expected, *Mimiviridae* was the family with dominant connections to virophages (Fig. 6A). Recently, *in silico* evidence demonstrated that virophages can infect *Phycodnaviridae*, which indicated that the virophage host range might be larger than we know (55). In support of this hypothesis, a relatively large number of virophage OTUs were found to be associated with *Phycodnaviridae* in our study. The enrichment analysis also revealed significant connections between three virophage clades and NCLDV families (Fig. 6B). To support the enrichment analysis, we conducted an HGT analysis because gene transfers have previously been found between Sputnik virophages and giant viruses (56). Our HGT analysis indicated a previously undescribed infectious relationship between virophage clade A and *Iridoviridae*. Overall, the results of virophage–NCLDV associations support our previous statement that co-occurrence networks inference and analysis are appropriate for investigating NCLDV interactions in marine metagenomic data.

## Materials and Methods

### Metagenomic and metabarcoding data

The microbial metagenomic and eukaryotic metabarcoding data used in this study were previously generated from plankton samples collected by the *Tara* Oceans expedition from 2008 to 2013 (57, 58). Because our research requires paired metagenomic and metabarcoding datasets, we used data derived from the euphotic zone samples, namely those from the surface (SRF) and Deep Chlorophyll Maximum (DCM) layers (59). Type B DNA polymerase (*polB*) was used as the marker gene for NCLDVs. A total of 6818 NCLDV *polB* OTUs were extracted from the metagenomic datasets (i.e., the second version of the Ocean Microbial Reference Gene Catalog, OM-RGC.v2) using the pplacer phylogenetic placement method (ML tree) (22, 60, 61). These *polB* sequences were classified into seven NCLDV families (*Mimiviridae, Phycodnaviridae, Marseilleviridae, Ascoviridae, Iridoviridae, Asfarviridae*, and *Poxviridae*) and two other giant virus groups (“Medusaviridae” and “Pithoviridae”). For eukaryotes, we employed used the metabarcoding data for eukaryotes, which targeting the 18S ribosomal RNA gene hypervariable V9 region (V9) (62). Taxonomic annotation of the eukaryotic metabarcoding data was previously performed by the *Tara* Oceans consortium using an extensive V9_PR2 reference database (59), which was derived from the original Protist Ribosomal Reference (PR2) database (63). The diversity index of eukaryotic communities was calculated using the package “vegan” (64). Processed frequency data are available from GenomeNet (ftp://ftp.genome.jp/pub/db/community/tara/Cooccurrence).

### Data processing

A relative abundance matrix for the NCLDV *polB* OTUs was extracted from OM-RGC.v2 for the samples derived from the pico-size fractions (0.22–1.6 or 0.22–3.0 µm). We converted the relative abundances of *polB* OTUs back to absolute read counts based on gene length and read length (assumed to be 100 nt). This process was required because small decimal numbers cannot be used by FlashWeave and because relative abundance data suffer from apparent correlations, which reduce the specificity of co-occurrence networks in revealing microbial interactions (44). To build comprehensive interaction networks involving eukaryotes of different sizes, we extracted the V9 read count matrices from the metabarcoding dataset for the following five size fractions: 0.8–5 μm; 5–20 μm and 3–20 μm (hereafter referred to as “5–20 μm” for simplicity); 20–180 μm; 180–2000 μm; and > 0.8 μm (hereafter referred to as 0.8–inf μm). To create the input files for network inference, the *polB* matrix was combined with each of the V9 matrices (corresponding to different size fractions), and only the samples represented by both *polB* and V9 files were placed in new files. In total, samples from 84 *Tara* Oceans stations (a total of 560 samples for two depths and five size fractions) widely distributed across oceans were used in this study (Fig. S1A). Depending on the individual size fractions, 84–127 samples were retained and included in the co-occurrence analysis (Fig. S1B). Read counts in the newly generated matrices were normalized using centered log-ratio (*clr*) transformation after adding a pseudo count of one to all matrix elements because zero cannot be transformed in *clr*. Following *clr* normalization, we filtered out low-abundance OTUs with a lower quartile (Q1) filtering approach. Specifically, OTUs were retained in the matrices when their *clr*-normalized abundance was higher than Q1 (among the non-zero counts in the original count matrix prior to the addition of a pseudo count of one) in at least five samples. Normalization and filtering were separately applied to *polB* and V9. The numbers of OTUs in the final matrices are provided in Fig. S1C.

### Co-occurrence-based network inference

Network inference was performed using FlashWeave [v0.15.0 (47)]. FlashWeave is a fast and compositionally robust tool for ecological network inference based on the local-to-global learning framework. Meta-variables (such as environmental parameters) can be included in the FlashWeave network to remove potential indirect associations. We used temperature, salinity, nitrate, phosphate, and silicate concentrations as meta-variables in our network inferences to determine their correlations with *polB* OTUs. FlashWeave provides a heterogeneous mode (FlashWeave-HE), which helps overcome sample heterogeneity. However, FlashWeave-HE may not be appropriate for the *Tara* Oceans data because it was shown to predict an insufficient number of known planktonic interactions (47). Therefore, we mainly used FlashWeave-S with default settings except for the FlashWeave normalization step and comparison between FlashWeave-S and FlashWeave-HE. A threshold to determine the statistical significance was set to alpha < 0.01. All detected pairwise associations were assigned a value called “weight” that ranged between −1 and +1. Edges with weights > 0 or < 0 were referred to as positive and negative associations, respectively. To compare the performance of FlashWeave-S to other co-occurrence methods, we used FlashWeave-HE, Spearman, and FastSpar (65). The FlashWeave-HE settings were the same as FlashWeave-S but with a command “heterogeneous”. For Spearman, we used stats.spearmanr in package “Scipy” (66). In FastSpar, we used 50 iterations, 20 excluded iterations, and a threshold of 0.1 to generate associations. To reduce the high dimensionality of the datasets, upper quartile (Q3) filtered matrices were used for comparison among FlashWeave-S, Spearman, and FastSpar.

### Network validation

We validated the virus–host associations in inferred networks based on a confusion matrix defined by the known NCLDV–host information (Fig. 1). Briefly, we manually compiled 69 known virus–host relationships for NCLDVs (Table S4). In the validation process, eukaryotic taxonomic groups were annotated at the level of the “Major lineages” in the extensive PR2 database (updated after publication) (62). The “Major lineages” were used in the present study because 1) the deficiency of known virus–host relationships limited the use of lower eukaryotic taxonomy ranks, such as genus, for assessment, and 2) these lineages adequately represented marine eukaryotes by covering the full spectrum of cataloged eukaryotic V9 diversity at a comparable phylogenetic depth (62). Then, we performed BLASTp [2.10.1 (67)] searches from the *Tara* Oceans PolB sequences against the NCLDV reference database to define groups of metagenomic PolBs with a threshold of 65% sequence identity by retaining only the best hit for each environmental PolB sequence. This threshold was determined because, by using reference PolB sequences and RefSeq protein sequence databases, we found that 60–70% of sequence identity could distinguish whether the NCLDVs infected hosts of the same major lineages; this was mainly tested for *Phycodnaviridae* because of the lack of host information for closely related viruses in other NCLDV families (Table S5). Then, 65% was chosen because it could provide a better LR+ (as described below) than 60% and 70%.

The positive likelihood ratio was used in for assessment to estimate the predictions accuracy. This approach is commonly used in diagnostic testing to assess if a test (host prediction in this study) usefully changes the probability of the existence of condition positive (true positive). In this study, the LR+ was used because host prediction is a test to discover condition positive states (68). LR+ is calculated by dividing the true-positive rate (sensitivity) by the false-positive rate (1 –specificity). If LR+ is close to 1, the performance of the prediction is comparable to a random prediction. If LR+ >> 1, a positive prediction result is more likely to be a true positive than that based on random prediction. From the set of detected associations between a given *polB* OTU and V9 OTUs that belong to a given major eukaryotic lineage, we only kept the best positive and negative associations (i.e., the edges with the highest absolute weights) to simplify the prediction scheme. As an auxiliary assessment, the FDR was also calculated by dividing the number of false positives by the number of positive predictions (Fig. 1). For the comparison among five size fractions, we only used the abundance in the overlap samples of 0.8–5 µm, 5–20 µm, 20–180 µm, and 180–2000 µm sizes. So the number of samples in five size fractions is comparable (84, 88, 88, 88, 88), which could reduce the bias that may influence the topology of networks (69).

### Phylogeny-guided filtering of host predictions and its assessment

We developed Taxon Interaction Mapper (TIM) to improve host predictions by co-occurrence approaches (33). TIM assumes that evolutionarily related viruses tend to infect evolutionarily related hosts (17, 70), and extract the most likely virus-host associations from the co-occurrence networks. TIM requires a phylogenetic tree of viruses (based on marker genes) and a set of connections between viruses and eukaryotes (co-occurrence edges), and then tests whether leaves (i.e., viral OTUs) under a node of the virus tree is enriched with a specific predicted host group compared with the rest of the tree using Fisher’s exact test and Benjamini–Hochberg adjustment (Fig. S9A) (33). TIM is available from https://github.com/RomainBlancMathieu/TIM.

We pooled network associations using FlashWeave analysis for five size fractions. To build a concise and credible viral phylogenetic tree, we removed all of the PolB sequences that were absent in the FlashWeave network associations, and the remaining sequences were filtered by the amino acid sequence length (≥ 500 aa). Protein alignment was conducted using MAFFT-*linsi* [version 7.471 (71)], and 18 sequences were manually removed because they were not well aligned with other PolB sequences. A total of 501 PolB sequences were used to make a maximum likelihood phylogenetic tree with FastTree [version 2.1.11 (72)]. Then, the PolB–V9 associations were mapped on the tree to calculate the significance of the enrichment of specific associations using TIM. TIM provides a list of nodes in the viral tree and associated NCBI taxonomies (order, class, and phylum) of eukaryotes that show significant enrichment in the leaves under the nodes. The TIM result was visualized with iTOL [version 5 (73)]. The TIM result was converted to a network, in which nodes correspond to the major eukaryotic lineages. The network edge weight was defined by the number of tree nodes in each viral family subtree enriched with a specific major eukaryotic lineage. The network was visualized with Cytoscape [version 3.7.1] using prefuse force directed layout (74). To assess the effectiveness of TIM in improving prediction, we extracted all the associations predicted by TIM and compared their performance with the raw and weight cut-off results.

### Virophage–NCLDV associations

We inferred the networks between NCLDVs and virophages using. *mcp* was used as the marker gene for virophages. First, 47 reference MCP amino acid sequences were collected from public databases and used to build an HMM profile. The HMM profile was used to search against the amino acid sequences of OM-RGC v2 using HMMER hmmsearch [version 3.3.1] with the threshold of E-value < 1E−90 (75). This threshold was determined by searching reference sequences against the GenomeNet nr-aa database. The search detected 195 *Tara* Oceans virophage MCP sequences in the OM-RGC database. Together with 47 reference MCPs, a phylogenetic tree of MCP amino acid sequences was built using MAFFT and FastTree.

We extracted the abundance profiles for the 195 MCP sequences from the pico- (0.22–1.6 or 0.22–3.0 µm) and femto-size (< 0.22 µm) fractions. We used samples from the SRF and DCM depths. PolB and MCP abundance profiles were merged into two matrices corresponding to the two virophage size fractions. Then, network inference was conducted using the FlashWeave default settings after Q1 filtration. In the MCP phylogenetic tree, three virophage clades contributed most of the NCLDV connections. Thus, an NCLDV enrichment analysis for the three clades was carried out using Fisher’s exact test, and the *p*-value was adjusted by the Benjamin–Hochberg method. This approach was the same as TIM, but we did not use the TIM software because the current version of TIM requires inputs of eukaryotic nodes with NCBI taxonomy annotations.

We used another approach, HGT, to predict the virophage-NCLDV interactions. First, we generated an NCLDV genome database, which includes 56 reference NCLDV genomes corresponding to our *polB* dataset and 2,074 metagenome-assembled genomes from a previous study (24). A total of 827,548 coding sequences were included in this database. Then, 195 virophage MCPs from the metagenomic data were BLASTp searched against this database using an E-value cut-off of 1E−10 (with a minimum query coverage of 50% and a minimum sequence identity of 50%). If a virophage MCP obtained a hit in the NCLDV genome database with a lower E-value compared with hits in the MCP database (the hit to itself was removed), the hit in the NCLDV genome database was considered an HGT candidate.

## Supporting information

all supplementaries

## Acknowledgments

This work was supported by JSPS/KAKENHI (Nos. 18H02279 and 19H05667 to H.O. and Nos. 19K15895 and 19H04263 to H.E.), Kyoto University Research Coordination Alliance (funding to H.E.), and the Collaborative Research Program of the Institute for Chemical Research, Kyoto University (Nos. 2019-30 and 2020-27). Computational time was provided by the SuperComputer System, Institute for Chemical Research, Kyoto University. We further thank the *Tara* Oceans consortium, and the people and sponsors who supported *Tara* Oceans. *Tara* Oceans (including both the *Tara* Oceans and *Tara* Oceans Polar Circle expeditions) would not exist without the leadership of the *Tara* Expeditions Foundation and the continuous support of 23 institutes (https://oceans.taraexpeditions.org). This article is contribution number XXX of *Tara* Oceans. We thank Mallory Eckstut, PhD, from Edanz Group (https://en-author-services.edanzgroup.com/ac) for editing a draft of this manuscript.

## Author contributions

LM and HO designed the study. LM performed most of the bioinformatics analysis.

HE generated the primary NCLDV data. RBM, SC, RHV, and HK contributed to the bioinformatics analysis. All authors contributed to the writing of the manuscript.

## Materials & Correspondence

Correspondence and material requests should be addressed to HO.

