## Supplementary material for "Quantitative assessment of NCLDV–host interactions predicted by co-occurrence analyses": all supplementaries

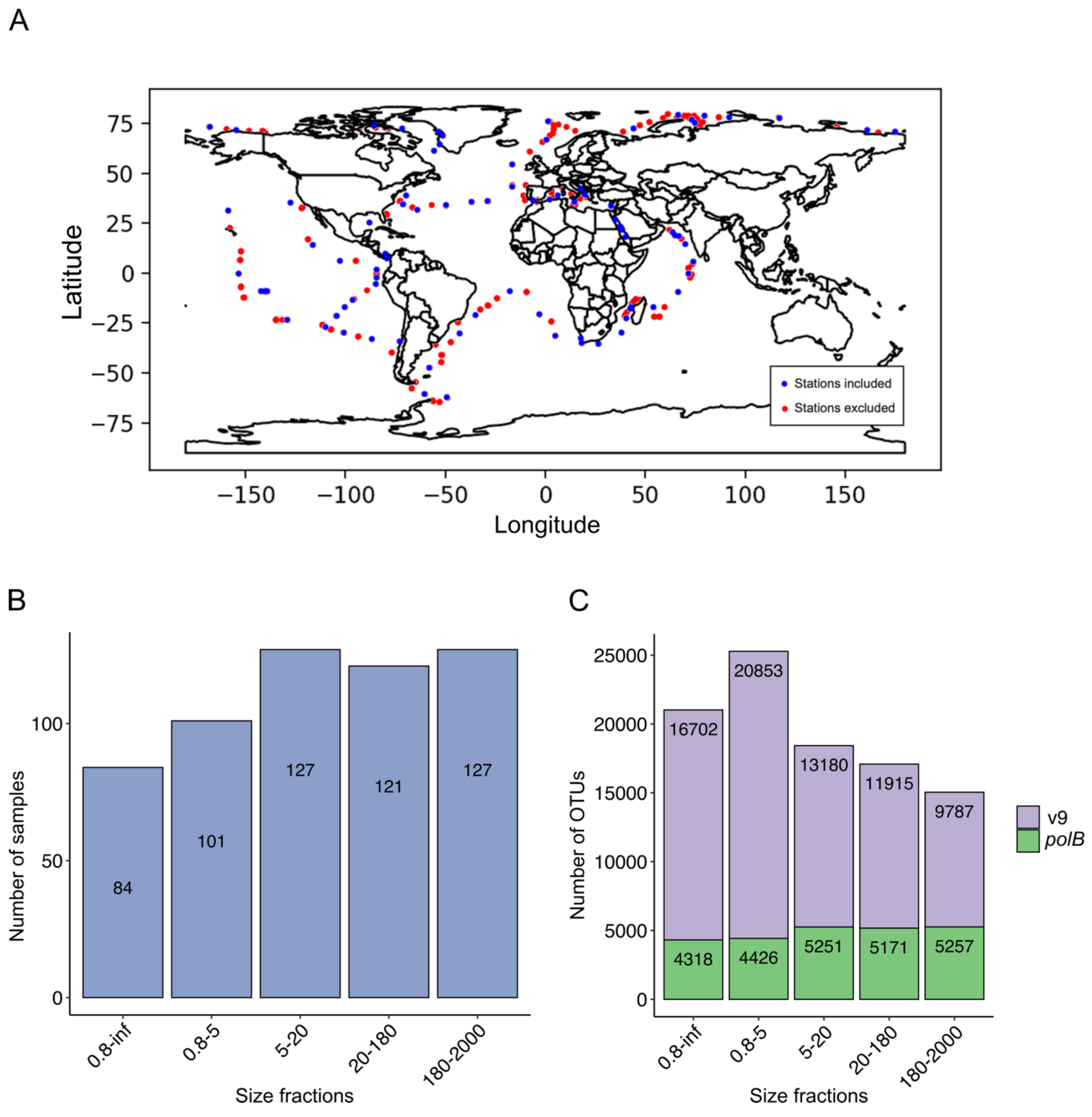


**Supplementary Figure 1. Stations, samples, and OTUs used in this study.** The *Tara* Oceans expedition is an international and multidisciplinary project; the stations are distributed worldwide. (A) After dataset treatment, the samples from a total of 84 stations were used in this study, and the retained stations were evenly distributed in the ocean. (B) Depending on the individual size fractions, 84–127 samples were retained and included in the co-occurrence analysis. (C) Numbers of OTUs (V9 and *polB* are presented seperately) in the final five individual size fraction matrices. Size fractions are presented in μm.

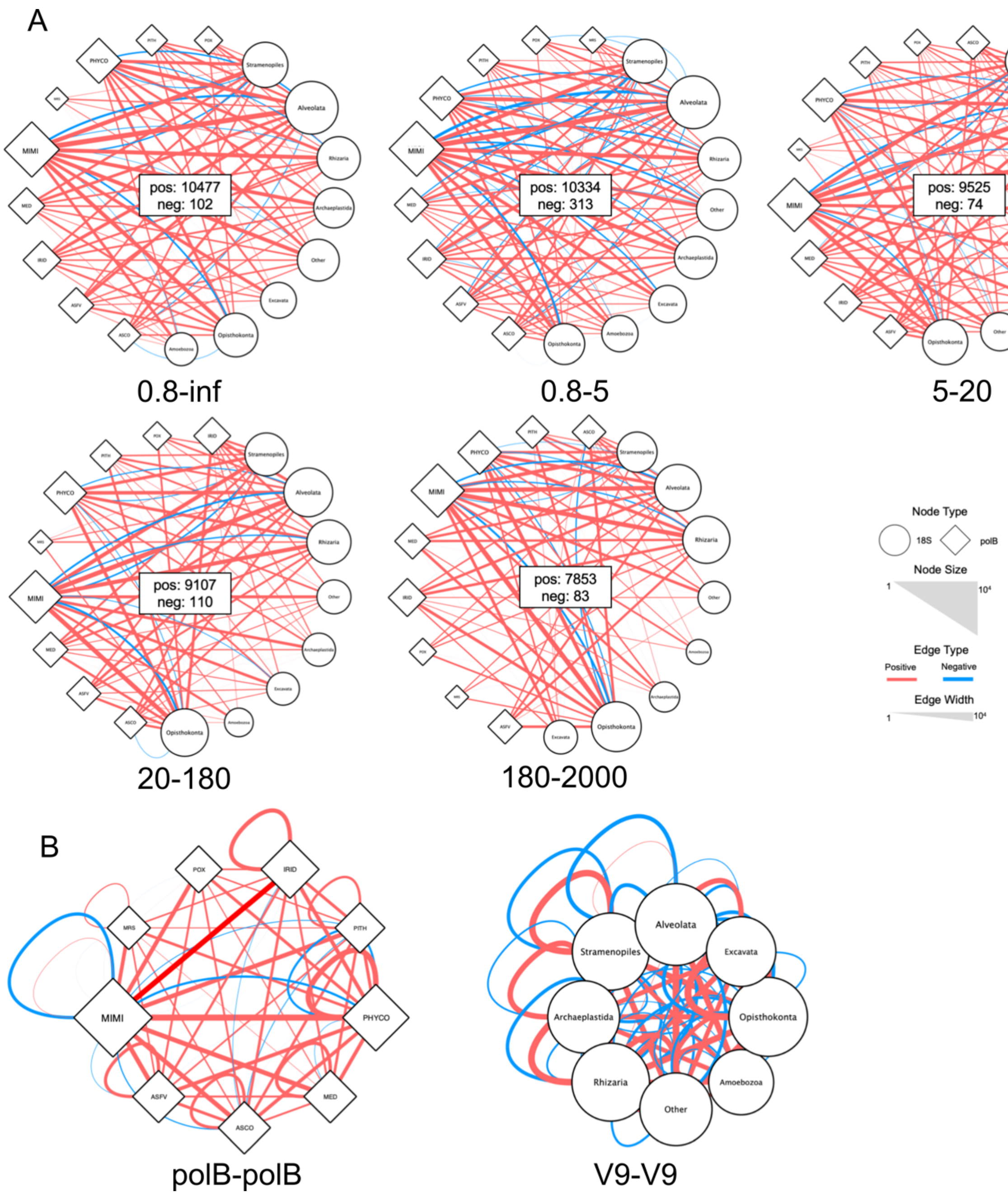


**Supplementary Figure 2. Co-occurrence networks inferred using FlashWeave.** The size of each node indicates the number of OTUs that belong to the group, and the width of each edge indicates the number of associations between two connected groups. Associations with positive weight are shown in red and negative associations are shown in blue. (A) ***polB*** –V9 co-occurrence networks of four individual size fractions and a broad range of size fractions (0.8–inf µm). (B) Networks consist of ***polB*** – ***polB*** and V9–V9 associations. Both associations were pooled from five size fraction networks. Size fractions are presented in µm.

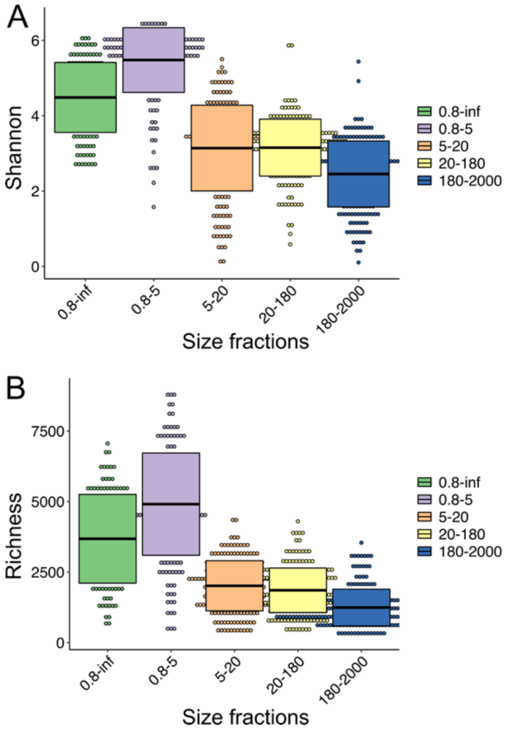


**Supplementary Figure 3. The diversity index of eukaryotic communities in five size fractions.** (A) Shannon diversity and (B) richness index was calculated for eukaryotic communities, from small piconanoplankton (0.8–5 μm) to large mesoplankton (180–2000 μm). Only the samples involved in co-occurrence inference for host prediction were used. Box plots are shown to summarize variation in richness and Shannon's index across (centerline, median; box limits, 25%–75% quantiles).

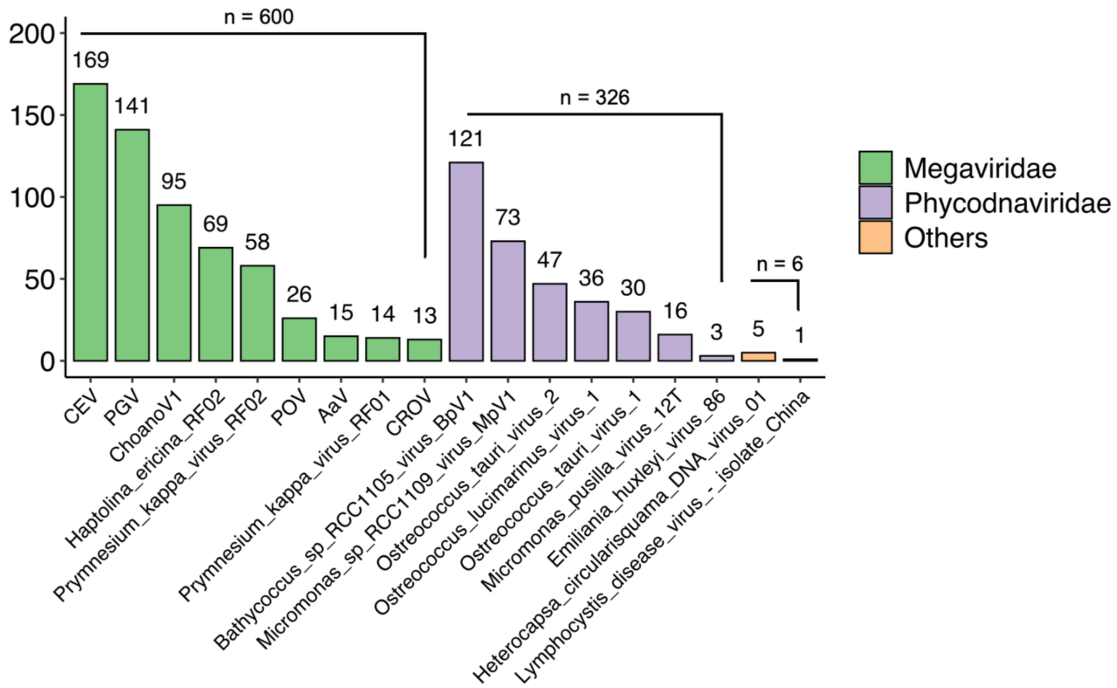


**Supplementary Figure 4. Number of environmental PolB OTUs recruited in the network validation.** The 932 OTUs were grouped using 69 reference NCLDV sequences. Apart from *Phycodnaviridae* and *Mimiviridae*, other families did not recruit many environmental OTUs; these families are shown together under an additional group, “Others.”

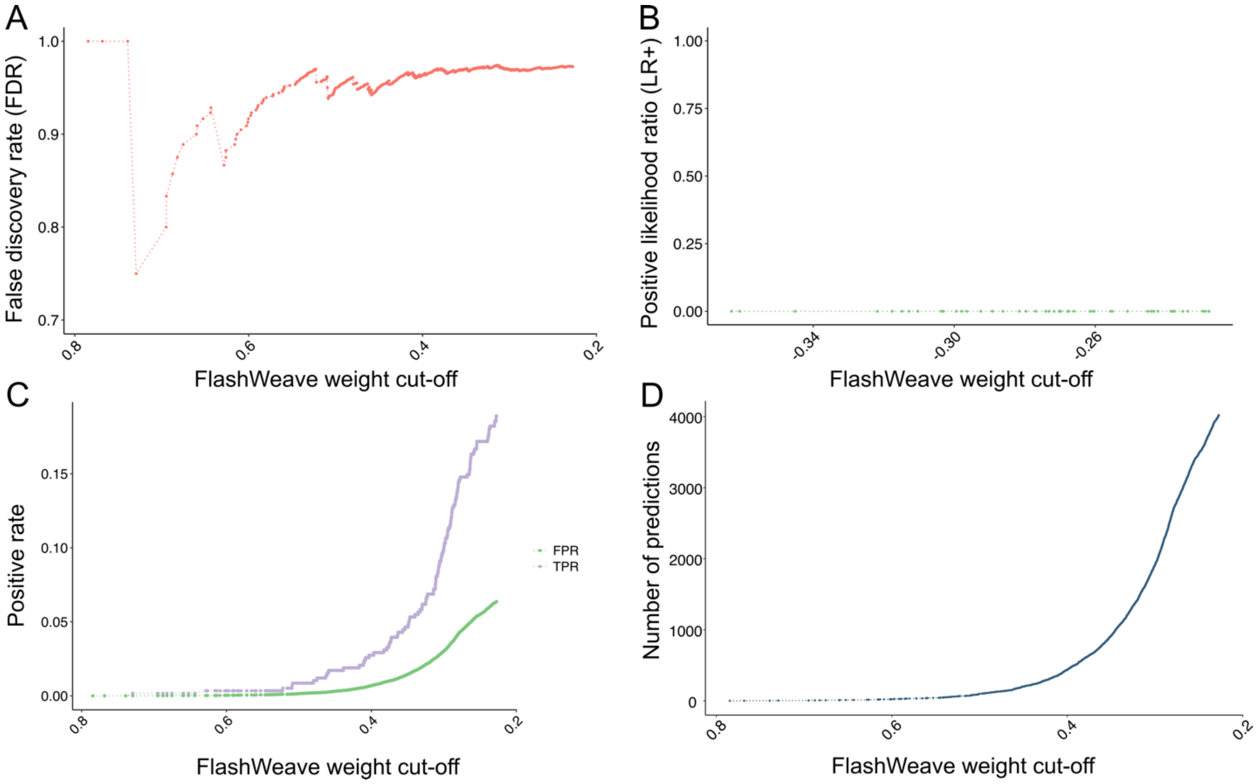


**Supplementary Figure 5. Performance of networks in the validation of NCLDV virus–host prediction.** (A) False discovery rate of the co-occurrence network, which consists of positive associations pooled from five size fractions networks. (B) Positive likelihood ratio (LR+) of the co-occurrence network, which consists of negative weight associations pooled from five size fractions networks. (C) Positive rate of the co-occurrence network, which consists of positive associations pooled from five size fractions networks. True positive rate (sensitivity, purple) and false positive rate (green) are given. (D) Number of predicted ***polB*** –V9 positive associations of the co-occurrence network.

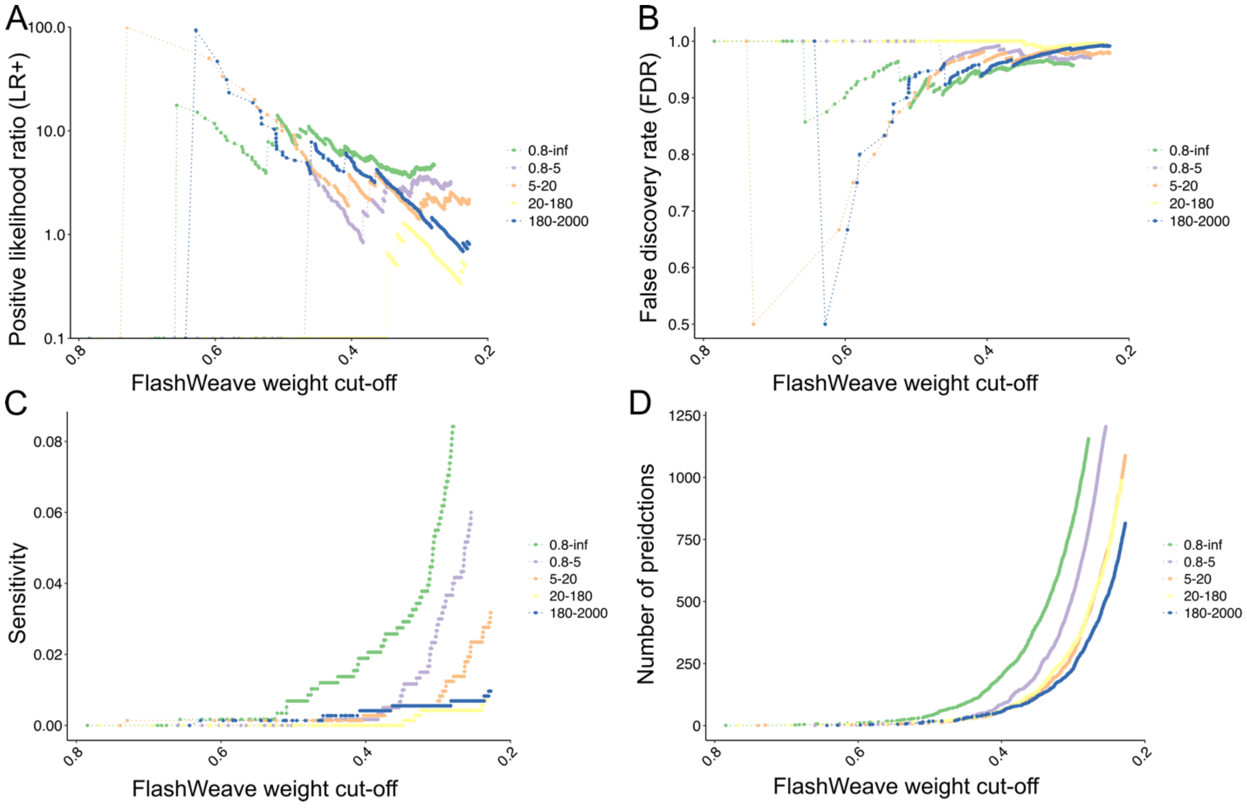


**Supplementary Figure 6. Comparison of different size fraction networks in the validation of NCLDV virus–host prediction.** (A) Positive likelihood ratio (LR+) of co-occurrence networks of five individual size fraction networks. (B) False discovery rate of co-occurrence networks of five individual size fractions networks. (C) Sensitivity of co-occurrence networks of five individual size fraction networks. (D) Number of predicted ***polB*** –V9 positive associations of co-occurrence networks of five individual size fraction networks. Size fractions are presented in µm.

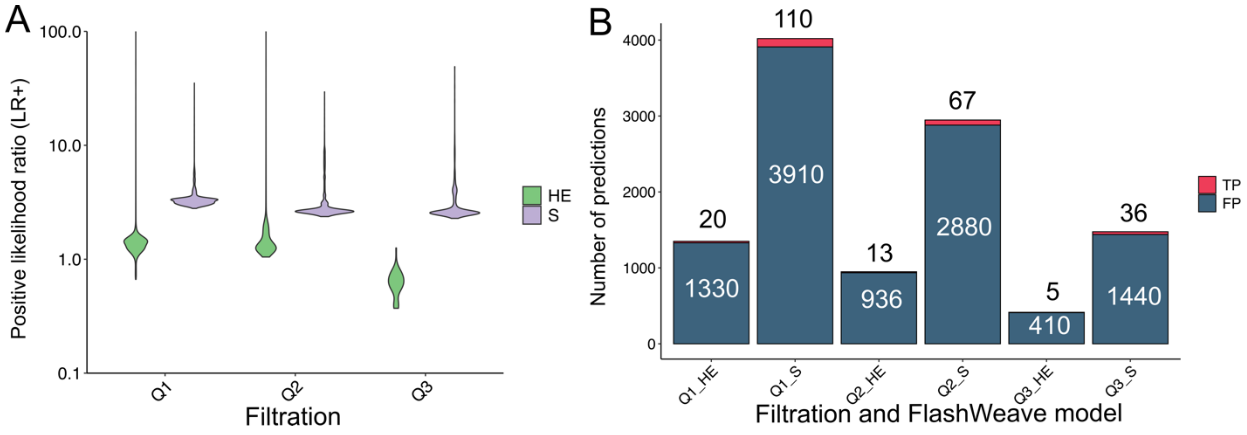


**Supplementary Figure 7. Performance of different input data trimming strategies.** To show if the OTU abundance influences the performance of the networks in the NCLDV virus–host validation, the datasets were treated by three quartile filtrations before the network inference, and the positive likelihood ratios (LR+) were compared. "S", FlashWeave sensitive model, and "HE", FlashWeave heterogeneous model, were performed separately. (A) Positive likelihood ratio (LR+) of networks inferred through different strategies. The plots are shown with a violin plot constructed by ggplot2 with a bandwidth of 2. (B) The number of inferred positive weight associations through different strategies, the number of ***polB*** –V9 pairs recruited in the validation is given

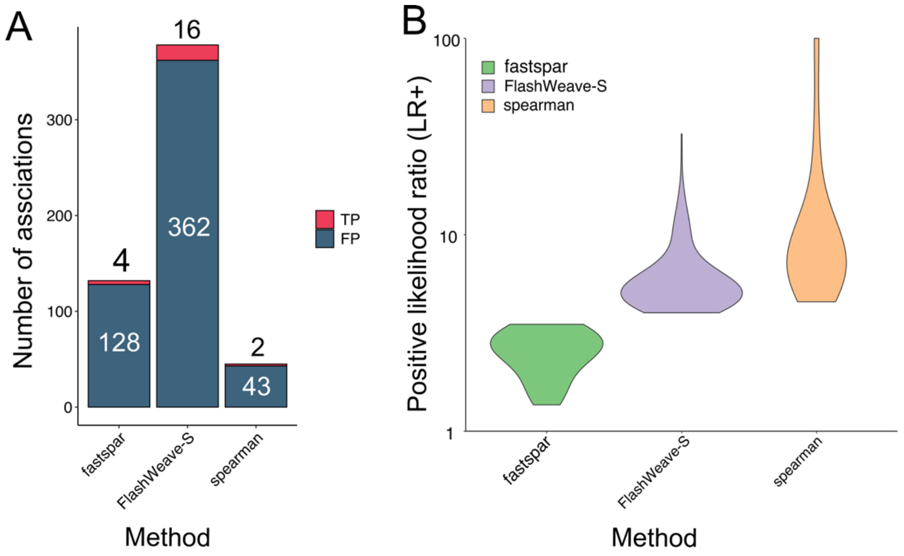


**Supplementary Figure 8. Performance of different network inference methods.** The networks were inferred using FastSpar, FlashWeave sensitive model, and spearman. (A) The number of inferred positive weight associations by three methods, the number of ***polB*** –V9 pairs recruited in the validation is given (true positive and false positive are shown separately). (B) The positive likelihood ratio (LR+) of the pooled associations of the five size fractions inferred by different methods were compared. To reduce the time of network inference in the large *Tara* Oceans datasets, upper quartile (Q3) trimming was performed on the input data.

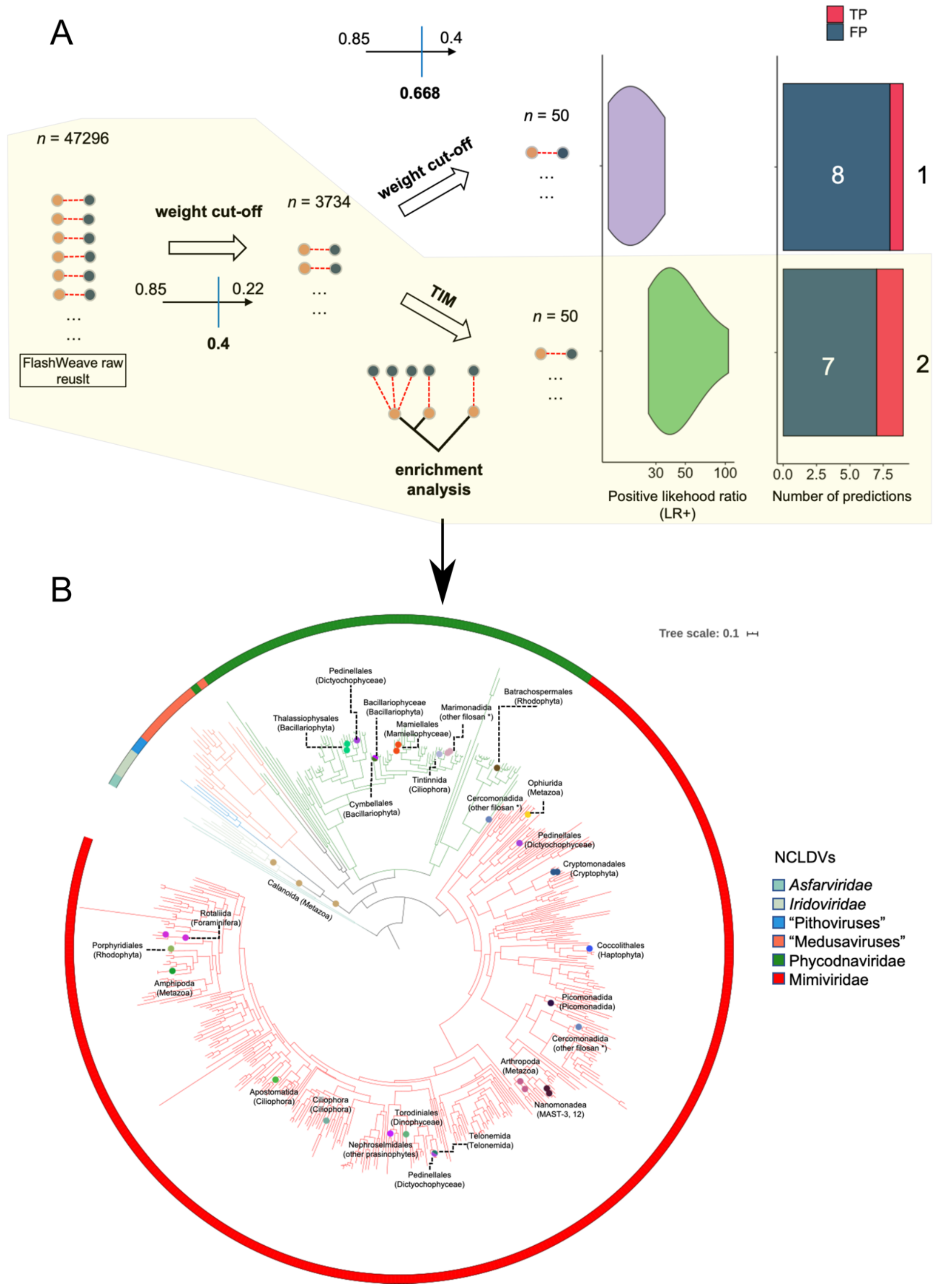


**Supplementary Figure 9. Filtration of FlashWeave results.** (A) The process of filtration. The number of retained *polB* –V9 pairs are given by *n*, we also compared the performance between TIM filtration and a further weight cut-off (0.668 was used because it provided the same number of *polB* –V9 pairs as TIM). The process used for making the final filtration and prediction is shown in yellow. (B) Phylogenetic position of NCLDVs and corresponding TIM-based predicted eukaryotic host groups. The phylogenetic tree was constructed from 501 *Tara* Oceans PolB protein sequences. Predicted hosts were shown with colored circles and annotated by NCBI taxonomies (order, class, and phylum).

#### Supplementary Table 1. Associations between NCLDVs and environmental variables inferred by FlashWeave.

| Size_fractions | PolB | Family | Envrionmental_parameter | Association_weight |
|---|---|---|---|---|
| 0.8-5 | TARA_138_DCM_0.22-3_G_C18424242_1_gene501927 | Ascoviridae | NO2 | 0.572042343 |
| 0.8-inf | TARA_180_SRF_lt-0.22_G_scaffold9939_5_gene48951 | Ascoviridae | Salinity | -0.294261709 |
| 20-180 | TARA_206_MES_lt-0.22_G_scaffold78978_1_gene218156 | Ascoviridae | Temperature | -0.273190139 |
| 0.8-5 | TARA_155_DCM_0.22-3_G_C16978172_1_gene567032 | Medusavirus | NO2 | 0.293908346 |
| 20-180 | TARA_007_SRF_0.22-1.6_G_scaffold79565_1_gene70629 | Megaviridae | Temperature | 0.300252947 |
| 5-20 | TARA_009_SRF_0.22-1.6_G_scaffold107825_1_gene135836 | Megaviridae | Temperature | 0.311369049 |
| 5-20 | TARA_031_SRF_0.22-1.6_G_C28711937_1_gene471690 | Megaviridae | Salinity | 0.24291636 |
| 20-180 | TARA_038_DCM_0.22-1.6_G_C23589122_1_gene515547 | Megaviridae | NO2 | 0.238884591 |
| 180-2000 | TARA_042_SRF_0.22-1.6_G_scaffold239468_1_gene192167 | Megaviridae | Temperature | 0.29866236 |
| 180-2000 | TARA_076_SRF_0.22-0.45_G_scaffold67722_1_gene45184 | Megaviridae | Temperature | 0.26981319 |
| 180-2000 | TARA_084_SRF_0.22-3_G_scaffold217252_1_gene156539 | Megaviridae | PO4 | 0.31500122 |
| 5-20 | TARA_085_DCM_0.22-3_G_scaffold263925_1_gene243728 | Megaviridae | Si | 0.260688454 |
| 0.8-5 | TARA_085_SRF_0.22-3_G_C16197045_1_gene301689 | Megaviridae | Temperature | -0.41365023 |
| 180-2000 | TARA_085_SRF_0.22-3_G_scaffold143071_1_gene112591 | Megaviridae | Si | 0.279735327 |
| 0.8-inf | TARA_102_DCM_0.22-3_G_scaffold400037_1_gene474877 | Megaviridae | NO2 | 0.279678103 |
| 180-2000 | TARA_111_SRF_0.22-3_G_scaffold275213_1_gene259640 | Megaviridae | NO2 | 0.272621244 |
| 20-180 | TARA_122_DCM_0.22-0.45_G_C14241023_1_gene864905 | Megaviridae | Temperature | 0.284428686 |
| 180-2000 | TARA_125_MIX_0.22-0.45_G_scaffold296071_1_gene285960 | Megaviridae | Temperature | 0.325251741 |
| 0.8-5 | TARA_133_SRF_0.22-3_G_C26534727_1_gene887577 | Megaviridae | Salinity | -0.318847308 |
| 180-2000 | TARA_158_SRF_0.22-3_G_scaffold245594_1_gene258773 | Megaviridae | Temperature | -0.294924596 |
| 5-20 | TARA_163_SRF_0.22-3_G_scaffold123902_2_gene136645 | Megaviridae | Temperature | -0.290310396 |
| 180-2000 | TARA_082_DCM_0.22-3_G_scaffold104031_1_gene99802 | Phycodnaviridae | Temperature | -0.303506051 |
| 180-2000 | TARA_193_SRF_0.22-3_G_scaffold165463_1_gene212660 | Phycodnaviridae | Salinity | -0.300940931 |
| 5-20 | TARA_194_DCM_lt-0.22_G_scaffold15132_1_gene46748 | Phycodnaviridae | Temperature | -0.238486803 |
| 0.8-5 | TARA_140_SRF_0.22-3_G_C20955321_1_gene443592 | Pithoviridae | Salinity | -0.265100268 |

#### Supplementary Table 2. Converting taxonomic annotation from TIM to the major eukaryotic lineages.

| Mimiviridae | | | Phycodnaviridae | | | Iridoviridae | | |
|---|---|---|---|---|---|---|---|---|
| NCBI taxonomy | Major lineage | Number of eniched braches | NCBI taxonomy | Major lineage | Number of eniched braches | NCBI taxonomy | Major lineage | Number of eniched braches |
| Ophiurida | Metazoa | 1 | Thalassiophysales | Bacillariophyta | 2 | Calanoida | Metazoa | 3 |
| Pedinellales | Dictyochophyceae | 1 | Pedinellales | Dictyochophyceae | 1 | | | |
| Cryptomonadales | Cryptophyta | 2 | Bacillariophyceae | Bacillariophyta | 1 | | | |
| Coccolithales | Haptophyta | 1 | Cymbellales | Bacillariophyta | 1 | | | |
| Picomonadida | Picomonadida | 1 | Mamiellales | Mamiellophyceae | 2 | | | |
| Cercomonadida | other filosan * | 1 | Tintinnida | Ciliophora | 1 | | | |
| Nanomonadea | MAST-3, 12 | 2 | Marimonadida | other filosan * | 2 | | | |
| Arthropoda | Metazoa | 2 | Batrachospermales | Rhodophyta | 1 | | | |
| Pedinellales | Dictyochophyceae | 1 | Cercomonadida | other filosan * | 1 | | | |
| Telonemida | Telonemida | 1 | | | | | | |
| Torodiniales | Dinophyceae | 1 | | | | | | |
| Nephroselmidales | other prasinophytes | 1 | | | | | | |
| Ciliophora | Ciliophora | 1 | | | | | | |
| Apostomatida | Ciliophora | 1 | | | | | | |
| Amphipoda | Metazoa | 1 | | | | | | |
| Porphyridiales | Rhodophyta | 1 | | | | | | |
| Rotaliida | Foraminifera | 2 | | | | | | |

### Supplementary Table 3. MCPs of virophage for NCLDV HGT candidates

<table>
<tr><th>virophage_MCP_ID</th><th>NCLDV_CDS_ID</th><th>pident</th><th>length</th><th>evalue</th><th>bitscore</th><th>qlen</th><th>slen</th><th>qcovs</th></tr>
<tr><td>TARA_194_SRF_lt-0.22_G_scaffold100579_1_gene186703</td><td>GVMAG-S-3300007345-62|3300007345.a:Ga0070752_1001124_10</td><td>63.519</td><td>466</td><td>0</td><td>570</td><td>464</td><td>569</td><td>100</td></tr>
<tr><td>TARA_205_MES_lt-0.22_G_scaffold373061_1_gene463550</td><td>GVMAG-S-3300007345-62|3300007345.a:Ga0070752_1001124_10</td><td>78.704</td><td>216</td><td>3.80E-122</td><td>360</td><td>217</td><td>569</td><td>99</td></tr>
</table>

### Supplementary Table 4. The 69 reference NCLDVs used in this study.

| Virus | Class | Order | Family | Genus | Species | Major_lineage | Reference |
|---|---|---|---|---|---|---|---|
| **MIMI_ChoanoV1** | Unknown | Choanoflagellatea | Unknown | Unknown | Unknown | Choanoflagellatea | https://www.ncbi.nlm.nih.gov/pubmed/?t |
| **MIMI_CEV** | Haptophyta | Prymnesiales | Prymnesiaceae | Haptolina | ericina | Haptophyta | https://www.ncbi.nlm.nih.gov/pmc/article |
| **MIMI_Haptolina_ericina_RF02** | Haptophyta | Prymnesiales | Prymnesiaceae | Haptolina | ericina | Haptophyta | https://www.ncbi.nlm.nih.gov/pubmed/?t |
| **MIMI_PGV** | Haptophyta | Phaeocystales | Phaeocystaceae | Phaeocystis | globosa | Haptophyta | https://www.ncbi.nlm.nih.gov/pubmed/?t |
| **MIMI_Prymnesium_kappa_virus_RF02** | Haptophyta | Prymnesiales | Prymnesiaceae | Prymnesium | kappa | Haptophyta | https://www.ncbi.nlm.nih.gov/pubmed/?t |
| **MIMI_TetV-1** | Chlorodendrophyce | Chlorodendrales | Chlorodendraceae | Tetraselmis | Unknown | other core chloroph | https://www.sciencedirect.com/science/a |
| **MIMI_Prymnesium_kappa_virus_RF01** | Haptophyta | Prymnesiales | Prymnesiaceae | Prymnesium | kappa | Haptophyta | https://www.ncbi.nlm.nih.gov/pubmed/?t |
| **MIMI_AaV** | Pelagophyceae | Pelagomonadales | Pelagomonadacea | Aureococcus | anophagefferens | Pelagophyceae | https://www.sciencedirect.com/science/a |
| **MIMI_CROV** | Unknown | Bicoecea | Cafeteriaceae | Cafeteria | roenbergensis | Bicoecea | https://www.ncbi.nlm.nih.gov/pubmed/?t |
| **PHYCO_Ostreococcus_lucimarinus_virus_1** | Mamiellophyceae | Mamiellales | Bathycoccaceae | Ostreococcus | lucimarinus | Mamiellophyceae | https://www.genome.jp/virushostdb/8801 |
| **PHYCO_Ostreococcus_tauri_virus_1** | Mamiellophyceae | Mamiellales | Bathycoccaceae | Ostreococcus | tauri | Mamiellophyceae | https://www.genome.jp/virushostdb/6429 |
| **PHYCO_Ostreococcus_tauri_virus_2** | Mamiellophyceae | Mamiellales | Bathycoccaceae | Ostreococcus | tauri | Mamiellophyceae | https://www.genome.jp/virushostdb/6964 |
| **PHYCO_Micromonas_sp_RCC1109_virus_MpV1** | Mamiellophyceae | Mamiellales | Mamiellaceae | Micromonas | pusilla | Mamiellophyceae | https://www.genome.jp/virushostdb/8801 |
| **PHYCO_Micromonas_pusilla_virus_12T** | Mamiellophyceae | Mamiellales | Mamiellaceae | Micromonas | pusilla | Mamiellophyceae | https://www.genome.jp/virushostdb/7552 |
| **PHYCO_Bathycoccus_sp_RCC1105_virus_BpV1** | Mamiellophyceae | Mamiellales | Bathycoccaceae | Bathycoccus | prasinos | Mamiellophyceae | https://www.genome.jp/virushostdb/8801 |
| **PHYCO_Paramecium_bursaria_Chlorella_virus_FR483** | Trebouxiophyceae | Chlorellales | Chlorellaceae | Micractinium | conductrix | Trebouxiophyceae | https://www.genome.jp/virushostdb/3997 |
| **PHYCO_Acanthocystis_turfacea_Chlorella_virus_1** | Trebouxiophyceae | Chlorellales | Chlorellaceae | Chlorella | heliozoae | Trebouxiophyceae | https://www.genome.jp/virushostdb/3220 |
| **PHYCO_Paramecium_bursaria_Chlorella_virus_1** | Trebouxiophyceae | Chlorellales | Chlorellaceae | Chlorella | Unknown | Trebouxiophyceae | https://www.genome.jp/virushostdb/1050 |
| **PHYCO_Feldmannia_species_virus** | Phaeophyceae | Ectocarpales | Acinetosporaceae | Feldmannia | Unknown | Phaeophyceae | https://www.genome.jp/virushostdb/3942 |
| **PHYCO_Ectocarpus_siliculosus_virus_1** | Phaeophyceae | Ectocarpalesn | Ectocarpaceae | Ectocarpus | siliculosus | Phaeophyceae | https://www.genome.jp/virushostdb/3766 |
| **PHYCO_Emiliania_huxleyi_virus_86** | Phaeophyceae | Isochrysidales | Noelaerhabdaceae | Emiliania | huxleyi | Haptophyta | https://www.genome.jp/virushostdb/1810 |
| **PHYCO_Pandoravirus_salinus** | Unknown | Centramoebida | Acanthamoebidae | Acanthamoeba | castellanii | other Lobosa | https://www.genome.jp/virushostdb/1349 |
| **PHYCO_Pandoravirus_dulcis** | Unknown | Centramoebida | Acanthamoebidae | Acanthamoeba | castellanii | other Lobosa | https://www.genome.jp/virushostdb/1349 |
| **PHYCO_Heterosigma_akashiwo_virus_01** | Raphidophyceae | Chattonellales | Chattonellaceae | Heterosigma | akashiwo | Raphidophyceae | https://www.genome.jp/virushostdb/9719 |
| **MIMI_OLPV1** | Unknown | Unknown | Unknown | Unknown | Unknown | Unknown | |
| **MIMI_OLPV2** | Unknown | Unknown | Unknown | Unknown | Unknown | Unknown | |
| **MIMI_POV** | Unknown | Unknown | Unknown | Unknown | Unknown | Unknown | |
| **MIMI_megavirus_chiliensis** | Unknown | Centramoebida | Acanthamoebidae | Acanthamoeba | castellanii | other Lobosa | https://www.genome.jp/virushostdb/1094 |
| **MIMI_megavirus_Iba** | Unknown | Unknown | Amoebidae | Amoeba | Unknown | Tubulinea | https://www.genome.jp/virushostdb/1235 |
| **MIMI_megavirus_courdo7** | Unknown | Unknown | Amoebidae | Amoeba | Unknown | Tubulinea | https://www.genome.jp/virushostdb/1235 |
| **MIMI_moumouvirus_ochan** | Unknown | Centramoebida | Acanthamoebidae | Acanthamoeba | polyphaga | other Lobosa | https://www.ncbi.nlm.nih.gov/pubmed/?t |
| **MIMI_moumouvirus_monve** | Unknown | Centramoebida | Acanthamoebidae | Acanthamoeba | polyphaga | other Lobosa | https://www.ncbi.nlm.nih.gov/pubmed/?t |
| **MIMI_moumouvirus_M10A** | Unknown | Centramoebida | Acanthamoebidae | Acanthamoeba | polyphaga | other Lobosa | https://www.ncbi.nlm.nih.gov/pubmed/?t |
| **MIMI_terra2** | Unknown | Centramoebida | Acanthamoebidae | Acanthamoeba | polyphaga | other Lobosa | https://www.genome.jp/virushostdb/1128 |
| **MIMI_apmv** | Unknown | Centramoebida | Acanthamoebidae | Acanthamoeba | polyphaga | other Lobosa | https://www.genome.jp/virushostdb/2120 |
| **MIMI_hirudovirus_Sangsue** | Unknown | Centramoebida | Acanthamoebidae | Acanthamoeba | polyphaga | other Lobosa | https://www.ncbi.nlm.nih.gov/pmc/article |
| **MIMI_pointerouge1** | Unknown | Centramoebida | Acanthamoebidae | Acanthamoeba | polyphaga | other Lobosa | https://journals.plos.org/plosone/article? |
| **MIMI_Tupanvirus_soda_lake** | Unknown | Unknown | Hartmannellidae | Vermamoeba | Unknown | Tubulinea | https://www.nature.com/articles/s41467- |
| **MIMI_Tupanvirus_deep_ocean** | Unknown | Centramoebida | Acanthamoebidae | Acanthamoeba | castellanii | other Lobosa | https://www.nature.com/articles/s41467- |
| **MIMI_Klosneuvirus_KNV1_1** | Unknown | Unknown | Unknown | Unknown | Unknown | Unknown | |
| **MIMI_Namao_virus** | Actinopteri | Acipenseriformes | Acipenseridae | Acipenser | fulvescens | Metazoa | https://pubs.er.usgs.gov/publication/702 |
| **MED_Medusavirus_Unagene_392** | Unknown | Centramoebida | Acanthamoebidae | Acanthamoeba | castellanii | other Lobosa | https://www.ncbi.nlm.nih.gov/pmc/article |
| **MIMI_Klosneuvirus_KNV1_2** | Unknown | Unknown | Unknown | Unknown | Unknown | Unknown | |
| **MRS_lausannevirus** | Unknown | Centramoebida | Acanthamoebidae | Acanthamoeba | Unknown | other Lobosa | https://www.genome.jp/virushostdb/9998 |
| **MRS_T19** | Unknown | Centramoebida | Acanthamoebidae | Acanthamoeba | Unknown | other Lobosa | https://www.genome.jp/virushostdb/6945 |
| **ASCO_Heliothis_virescens_ascovirus_3e** | Insecta | Lepidoptera | Noctuidae | Heliothis | Unknown | Metazoa | https://www.genome.jp/virushostdb/2607 |
| **ASCO_Spodoptera_frugiperda_ascovirus_1a** | Insecta | Lepidoptera | Noctuidae | Spodoptera | Unknown | Metazoa | https://www.genome.jp/virushostdb/1133 |
| **ASCO_Trichoplusia_ni_ascovirus_2c** | Insecta | Lepidoptera | Noctuidae | Trichoplusia | Unknown | Metazoa | https://www.genome.jp/virushostdb/3286 |
| **IRID_Infectious_spleen_and_kidney_necrosis_virus** | Actinopteri | Syngnathiformes | Callionymidae | Synchiropus | Unknown | Metazoa | https://www.genome.jp/virushostdb/1801 |
| **IRID_Lymphocystis_disease_virus_-_isolate_China** | Actinopteri | Centrarchiformes | Centrarchoidei | Unknown | Unknown | Metazoa | https://www.genome.jp/virushostdb/3636 |
| **IRID_Lymphocystis_disease_virus_1** | Actinopteri | Centrarchiformes | Centrarchoidei | Unknown | Unknown | Metazoa | https://www.genome.jp/virushostdb/3636 |
| **IRID_Singapore_grouper_iridovirus** | Actinopteri | Perciformes | Serranidae | Epinephelus | Unknown | Metazoa | https://www.genome.jp/virushostdb/2629 |
| **IRID_Invertebrate_iridescent_virus_3** | Insecta | Diptera | Culicidae | Psorophora | Unknown | Metazoa | https://www.genome.jp/virushostdb/3452 |
| **IRID_Invertebrate_iridescent_virus_6** | Insecta | Orthoptera | Gryllidae | Gryllus | Unknown | Metazoa | https://www.genome.jp/virushostdb/1766 |
| **PITH_pithovirus** | Unknown | Centramoebida | Acanthamoebidae | Acanthamoeba | polyphaga | other Lobosa | https://www.genome.jp/virushostdb/1450 |
| **POX_Amsacta_moorei_entomopoxvirus_L** | Insecta | Lepidoptera | Lymantriidae | Lymantria | Unknown | Metazoa | https://www.genome.jp/virushostdb/2832 |
| **POX_Melanoplus_sanguinipes_entomopoxvirus** | Insecta | Orthoptera | Acrididae | Locusta | Unknown | Metazoa | https://www.genome.jp/virushostdb/8319 |
| **POX_Deerpox_virus_W-1170-84** | Mammalia | Cetartiodactyla | cervidae | Odocoileus | Unknown | Metazoa | https://www.genome.jp/virushostdb/3056 |
| **POX_Yaba-like_disease_virus** | Mammalia | Primates | Hominidae | Homo | Unknown | Metazoa | https://www.genome.jp/virushostdb/1324 |
| **POX_Monkeypox_virus_Zaire-96-I-16** | Mammalia | Primates | Hominidae | Homo | Unknown | Metazoa | https://www.genome.jp/virushostdb/6195 |
| **POX_Molluscum_contagiosum_virus_subtype_1** | Mammalia | Primates | Hominidae | Homo | Unknown | Metazoa | https://www.genome.jp/virushostdb/1028 |
| **POX_Orf_virus** | Mammalia | Primates | Hominidae | Homo | Unknown | Metazoa | https://www.genome.jp/virushostdb/1025 |
| **POX_Fowlpox_virus** | Aves | Galliformes | Phasianidae | Meleagris | Unknown | Metazoa | https://www.genome.jp/virushostdb/1026 |
| **POX_Nile_crocodilepox_virus** | Unknown | Crocodylia | Crocodylidae | Crocodylus | Unknown | Metazoa | https://www.genome.jp/virushostdb/1285 |
| **ASFV_African_swine_fever_virus** | Mammalia | Cetartiodactyla | Suidae | Sus | Unknown | Metazoa | https://www.genome.jp/virushostdb/1049 |
| **ASFV_Pacmanvirus_A23** | Unknown | Centramoebida | Acanthamoebidae | Acanthamoeba | Unknown | other Lobosa | https://www.ncbi.nlm.nih.gov/pubmed/28 |
| **ASFV_Heterocapsa_circularisquama_DNA_virus_01** | Dinophyceae | Peridiniales | Heterocapsaceae | Heterocapsa | Unknown | Dinophyceae | https://www.genome.jp/virushostdb/6501 |
| **MIMI_Bodo_saltans_virus** | Unknown | Kinetoplastida | Bodonidae | Bodo | Unknown | Kinetoplastida | https://elifesciences.org/articles/33014 |

**Notes:**

1.Cells filled with green:
  Viruses infect non-Amoeba protists

2.Major_lineage:
  85 deep-branching eukaryotic lineages represented in our global oceans metabarcodingdata set
  Reference:
  https://www.ncbi.nlm.nih.gov/pubmed/25999516

3.Some viruses, for example IRID_Lymphocystis_disease_virus_1, could infect more than one host species. But their hosts, large size metazoan animals, are not included in *Tara Oceans* sampling. So in this table, we only list one of their known hosts, more information could be found by reference links.

### Supplementary Table 5. Threshold determination of environmental NCLDV grouping.

| Ref_ID | The least hit with same host major lineage | Identity | Major Lineage | Accession | The best hit with different host major lineage | Identity | Major Lineage | Accession |
|---|---|---|---|---|---|---|---|---|
| **MIMI_ChoanoV1** | MIMI_ChoanoV1 | 100 | Choanoflagellatea | NULL | Acanthamoeba polyphaga mimivirus | 46.5 | Unknown | YP_003986825.1 |
| **MIMI_CEV** | MIMI_CEV | 100 | Haptophyta | NULL | Acanthamoeba polyphaga mimivirus | 40.47 | Unknown | YP_003986825.1 |
| **MIMI_Haptolina_ericina_RF02** | MIMI_CEV | 44.4 | Haptophyta | NULL | Bombus impatiens | 34.55 | Unknown | XP_003484917.1 |
| **MIMI_PGV** | MIMI_CEV | 44.96 | Haptophyta | NULL | Cafeteria roenbergensis virus BV-PW1 | 34 | Bicoecea | YP_003970130.1 |
| **MIMI_Prymnesium_kappa_virus_RF02** | Chrysochromulina ericina virus | 44.69 | Haptophyta | YP_009173620.1 | Entamoeba invadens IP1 | 34.65 | Unknown | XP_004185906.1 |
| **MIMI_TetV-1** | MIMI_TetV-1 | 100 | other core chlorophytes | NULL | MIMI_POV | 37.81 | Pyramimonadales | NULL |
| **MIMI_Prymnesium_kappa_virus_RF01** | Chrysochromulina ericina virus | 39.5 | Haptophyta | YP_009173620.1 | Cafeteria roenbergensis virus BV-PW1 | 36.76 | Bicoecea | YP_003970130.1 |
| **MIMI_AaV** | MIMI_AaV | 100 | Pelagophyceae | NULL | Acanthamoeba polyphaga mimivirus | 45.63 | other Lobosa | YP_003986825.1 |
| **MIMI_CROV** | Cafeteria roenbergensis virus BV-PW1 | 100 | Bicoecea | YP_003970130.1 | Megavirus chiliensis | 34.67 | other Lobosa | YP_004894633.1 |
| **PHYCO_Ostreococcus_lucimarinus_virus_1** | Micromonas pusilla virus 12T | 72.91 | Mamiellophyceae | YP_007676285.1 | Dishui lake phycodnavirus 1 | 59.6 | Unknown | YP_009465906.1 |
| **PHYCO_Ostreococcus_tauri_virus_1** | Bathycoccus sp. RCC1105 virus BpV1 | 74.37 | Mamiellophyceae | YP_004061614.1 | Dishui lake phycodnavirus 1 | 58.1 | Unknown | YP_009465906.1 |
| **PHYCO_Ostreococcus_tauri_virus_2** | Micromonas pusilla virus 12T | 73.2 | Mamiellophyceae | YP_007676285.1 | Dishui lake phycodnavirus 1 | 57.65 | Unknown | YP_009465906.1 |
| **PHYCO_Micromonas_sp_RCC1109_virus_MpV1** | Micromonas pusilla virus 12T | 71.71 | Mamiellophyceae | YP_007676285.1 | Dishui lake phycodnavirus 1 | 59.21 | Unknown | YP_009465906.1 |
| **PHYCO_Micromonas_pusilla_virus_12T** | Bathycoccus sp. RCC1105 virus BpV1 | 70.13 | Mamiellophyceae | YP_004061614.1 | Dishui lake phycodnavirus 1 | 56.99 | Unknown | YP_009465906.1 |
| **PHYCO_Paramecium_bursaria_Chlorella_virus_FR483** | Only Syngen Nebraska virus 5 | 65.98 | Trebouxiophyceae | YP_009325633.1 | Yellowstone lake phycodnavirus 1 | 38.68 | Unknown | YP_009174732.1 |
| **PHYCO_Acanthocystis_turfacea_Chlorella_virus_1** | Only Syngen Nebraska virus 5 | 63.97 | Trebouxiophyceae | YP_001427279.1 | Yellowstone lake phycodnavirus 1 | 39.04 | Unknown | YP_009174732.1 |
| **PHYCO_Paramecium_bursaria_Chlorella_virus_1** | Acanthocystis turfacea chlorella virus 1 | 71.9 | Trebouxiophyceae | YP_001427279.1 | Bathycoccus sp. RCC1105 virus BpV1 | 40.16 | Mamiellophyceae | YP_004061614.1 |
| **PHYCO_Feldmannia_species_virus** | Ectocarpus siliculosus virus 1 | 45.66 | Phaeophyceae | NP_077578.1 | Chlamydomonas reinhardtii | 34.57 | Unknown | XP_001689909.1 |
| **PHYCO_Ectocarpus_siliculosus_virus_1** | Feldmannia irregularis virus a | 44.16 | Phaeophyceae | YP_009665694.1 | Aedes aegypti | 32.28 | Unknown | XP_021705412.1 |
| **PHYCO_Emiliania_huxleyi_virus_86** | Emiliania huxleyi virus 86 | 100 | Haptophyta | YP_293784.1 | Agaricus bisporus var. burnettii JB137-S8 | 37.81 | Unknown | XP_007327213.1 |
| **PHYCO_Heterosigma_akashiwo_virus_01** | Heterosigma akashiwo virus 01 | 99.91 | Raphidophyceae | YP_009507574.1 | Yellowstone lake phycodnavirus 2 | 40.34 | Unknown | YP_009174650.1 |
| **ASFV_Heterocapsa_circularisquama_DNA_virus_01** | Heterocapsa circularisquama DNA virus 01 | 100 | Dinophyceae | YP_009507841.1 | Rhizophagus irregularis DAOM 181602=DAOM 19 | 26.03 | Unknown | XP_025181784.1 |
| **MIMI_Bodo_saltans_virus** | MIMI_Bodo_saltans_virus | 100 | Kinetoplastida | NULL | Megavirus chiliensis | 35.27 | other Lobosa | YP_004894633.1 |
| **MIMI_POV** | MIMI_POV | 100 | Pyramimonadales | NULL | MIMI_Prymnesium_kappa_virus_RF01 | 45.7 | Haptophyta | NULL |
